# An acidic residue buried in the dimer interface of isocitrate dehydrogenase 1 (IDH1) helps regulate catalysis and pH sensitivity

**DOI:** 10.1101/2020.04.19.049387

**Authors:** Lucas A. Luna, Zachary Lesecq, Katharine A. White, An Hoang, David A. Scott, Olga Zagnitko, Andrey A. Bobkov, Diane L. Barber, Jamie M. Schiffer, Daniel G. Isom, Christal D. Sohl

**Affiliations:** Department of Chemistry and Biochemistry, San Diego State University, San Diego, CA, 92182; Harper Cancer Research Institute, Department of Chemistry and Biochemistry, University of Notre Dame, South Bend, IN, 46617; Sanford Burnham Prebys Medical Discovery Institute, La Jolla, CA, 92037; Department of Cell and Tissue Biology, University of California, San Francisco, CA, 94143; Janssen Research and Development, 3210 Merryfield Row, San Diego, CA, 92121; Department of Pharmacology, Sylvester Comprehensive Cancer Center, and Center for Computational Sciences, University of Miami, Miami, FL, 33136

**Author notes:** To whom correspondence should be addressed: Christal D. Sohl, Department of Chemistry and Biochemistry, San Diego State University, CSL 328, 5500 Campanile Dr., San Diego, California 92182. This paper is dedicated to the memory of our dear colleague and friend, Michelle Evon Scott (1990-2020).

**Keywords:** enzyme kinetics, cancer, tumor metabolism, pH regulation, post-translational modification (PTM), buried ionizable residues

## Abstract

Isocitrate dehydrogenase 1 (IDH1) catalyzes the reversible NADP^+^-dependent conversion of isocitrate to α-ketoglutarate (α-KG) to provide critical cytosolic substrates and drive NADPH-dependent reactions like lipid biosynthesis and glutathione regeneration. In biochemical studies, the forward reaction is studied at neutral pH, while the reverse reaction is typically characterized in more acidic buffers. This led us to question whether IDH1 catalysis is pH-regulated, which would have functional implications under conditions that alter cellular pH, like apoptosis, hypoxia, cancer, and neurodegenerative diseases. Here, we show evidence of catalytic regulation of IDH1 by pH, identifying a trend of increasing *k*_cat_ values for α-KG production upon increasing pH in the buffers we tested. To understand the molecular determinants of IDH1 pH sensitivity, we used the pHinder algorithm to identify buried ionizable residues predicted to have shifted pK_a_ values. Such residues can serve as pH sensors, with changes in protonation states leading to conformational changes that regulate catalysis. We identified an acidic residue buried at the IDH1 dimer interface, D273, with a predicted pK_a_ value upshifted into the physiological range. D273 point mutations had decreased catalytic efficiency and, importantly, loss of pH-regulated catalysis. Based on these findings, we conclude that IDH1 activity is regulated, at least in part, by pH. We show this regulation is mediated by at least one buried acidic residue ∼12 Å from the IDH1 active site. By establishing mechanisms of regulation of this well-conserved enzyme, we highlight catalytic features that may be susceptible to pH changes caused by cell stress and disease.

## INTRODUCTION

Isocitrate dehydrogenase 1 (IDH1) is a highly conserved metabolic enzyme that catalyzes the reversible NADP^+^-dependent conversion of isocitrate to α-ketoglutarate (αKG) (Fig. 1A). This reversible reaction is important for providing substrates for a host of cytosolic reactions, for anaplerosis, or the restocking of the tricarboxylic acid (TCA) cycle, for providing reducing power in the form of NADPH, and for driving lipid metabolism (1-3).

**Figure 1.**
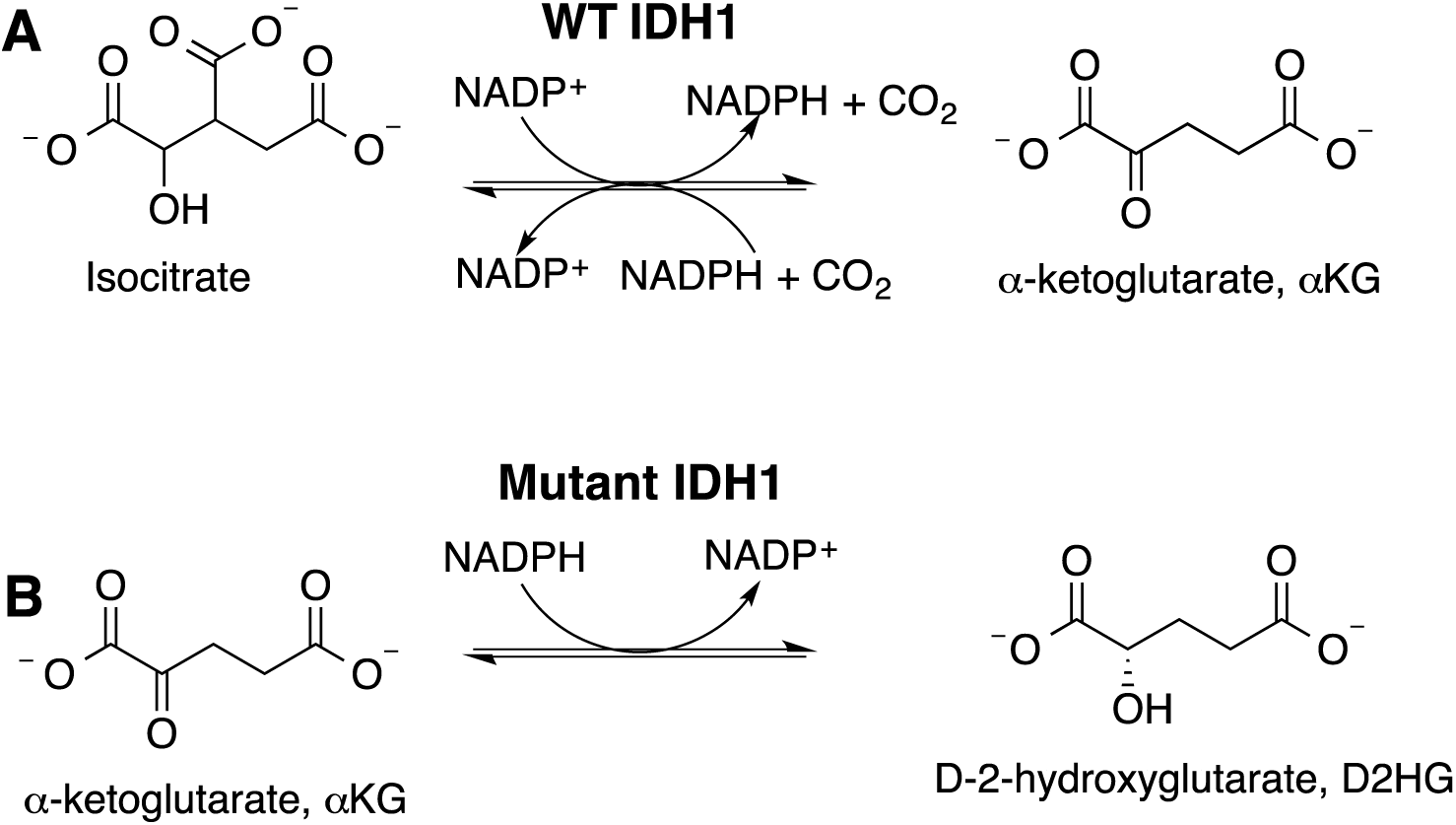
The normal and neomorphic reactions catalyzed by WT (A) and mutant (B) IDH1.

Given its importance in cell metabolism, it is perhaps unsurprising that changes in wild type (WT) IDH1 expression levels or acquisition of point mutations are important features of some cancers and result in interesting metabolic changes. For example, increased mRNA and protein levels of WT IDH1 are associated with cancer (4-6), including as high as 65% percent of primary glioblastomas (4). The most famous role of IDH1 in cancer is via the point mutations that drive ∼85% of lower grade gliomas and secondary glioblastomas, ∼12% of acute myeloid leukemias, and ∼40% of chondrosarcomas (7-11). Most mutations affect R132, an active site residue that plays a role in coordinating the C-3-carboxylate of isocitrate (12), and R132H and R132C are by far the most common mutations in IDH1 (5). These mutations confer an inability to catalyze the normal reaction, but facilitate a neomorphic reaction, the NADPH-dependent conversion of αKG to *D*-2-hydroxyglutarate (D2HG) (13) (Fig. 1B), an oncometabolite that inhibits αKG-binding enzymes (14,15). IDH1 mutants are bona fide therapeutic targets, with the first selective mutant IDH1 inhibitor recently approved for use in the clinic primarily for leukemias (16,17).

Proteins can be regulated by their environment to respond in real time to cellular needs. In response to stressors such as hypoxia, the reverse reaction catalyzed by IDH1, the reduction of αKG to isocitrate (Fig. 1A) appears to confer resilience (18-20). However, the mechanism of this metabolic shift is still under debate and may result from slowing of the forward reaction (21). It also remains unclear whether IDH1 (the cytosolic isoform) or IDH2 (the mitochondrial isoform) is the major player in reductive metabolism, with isoform localization, local substrate concentrations, and various mechanisms of regulation all likely complicating the discernment of the relative contributions of IDH1 and IDH2 (19,20,22-24).

Cellular pH plays a critical role in regulating protein-protein and protein-ligand interactions, as well as protein stability and activity (25). Unlike most protein regulatory mechanisms and post-translational modifications (PTMs), protonation is non-enzymatic, does not require ATP, and is rapidly reversible. As such, many cellular processes and human pathologies are regulated by small but meaningful changes in the intracellular pH (pH_i_) that are sufficient to alter residue ionization state and protein function. A variety of physiological and pathological cellular processes can cause these shifts in pH. For example, a decrease in pH_i_ can occur during apoptosis (26), immune processes (27), nutrient deprivation (28,29), and oxidative stress (30,31), while an increase in pH_i_ is important for cell differentiation (32) and directed cell migration (33). A decrease in pH_i_ is also associated with neurodegenerative diseases (34,35), while an increase in pH_i_ coupled with a decrease in extracellular pH (pH_e_) occurs in tumors to drive migration and metastasis (25,36-39).

Cellular proteins that can function as pH sensors detect changes in pH using ionizable amino acid residues. These residues sense changes in surrounding pH_i_ via changes in residue protonation/deprotonation, leading to biologically relevant changes in protein conformation. Proteins with pH sensitive residue(s) are known as pH sensors. H residues are natural candidates for sensing pH_i_ changes as their pK_a_ value is already in a physiologically relevant range. However, any ionizable residue (such as D, E, or K) can sense changes in pH if its pK_a_ value is shifted into the physiological pH range by protein structure (25,40). In fact, large changes in pK_a_ values, ΔpK_a_, make for stronger pH sensors. The change in Gibbs free energy difference, ΔΔG°, that may be stored within a change in pK_a_ value, ΔpK_a_, can be as high as ΔΔG° = 1.36 × Δp*K*_a_ (41). Interestingly, simply having a buried K residue in a protein tends to downshift that residue’s pK_a_ to a more physiological range. This may trigger conformational changes as a means of pH-regulated catalysis, or can simply be a mechanism to tune protein stability. As an example of the latter, variants of a staphylococcal nuclease were designed to test the consequences of buried K residues in a model protein system (42). The majority of the K variants show downshifted pK_a_ values to a more physiological range, with the range of shifted pK_a_ values varying widely (from 5 to 10) (42). The location of the lysine mutation appears to affect the degree of pK_a_ value shift due to changes in local environment; for example, interactions with carboxylic acid groups enhanced K pK_a_ value shifts (42). An extension of this work explores the consequences of comparing K, D, and E mutational variants at a single residue, with each variant having notable changes in pK_a_ values, protein stability, ionizability, and structural conformation (40). Thus, buried ionizable residues can play a critical role in both protein function, stability, and pH sensing.

There is significant diversity among the types of proteins that have been identified as pH sensors, including low molecular weight GTPases (43), G protein α subunits (29), G protein-coupled receptors (44), an Na^+^-H^+^ exchanger (45), β-catenin (46), a kinase (47), a guanine nucleotide exchange factor (48), and metabolic enzymes (49-57). Notable examples of metabolic enzymes include lactate dehydrogenase, whose activity is dependent on pH in part through changes in H residue protonation state that affects substrate binding and catalysis (58,59), and phosphofructokinase, which exhibits decreased catalytic efficiency driven primarily through an increase in *K*_m_ as the pH drops (29,55,60). Despite pH-sensing in phosphofructokinase being first described over half a century ago, the mechanisms of pH-sensitive catalysis are still not yet understood. Thus, identifying mechanisms of pH-sensing are challenging, but elucidating such mechanisms is potentially highly transformative for understanding environment-sensitive protein regulation.

There is evidence that IDH1 catalysis is also affected by changes in pH. Rates of NADPH consumption stemming from isocitrate production (the normal reverse reaction, Fig. 1A) by human WT IDH1 increase upon decreasing pH in potassium phosphate (KPhos) (61) and tris/bis-tris (62) buffers, though the effects on steady-state kinetic parameters are not reported. Pig IDH2 is sensitive to changes in pH, with increased rates of αKG production observed as pH increases (63). This appears to be driven in part by ionizability of H319 and H315 IDH2, as H319Q and H315Q (corresponding to H354 and H358 in human IDH2) decreases catalytic efficiency by ∼2-fold (63). However, only the residue corresponding to H354 in IDH2 is conserved in human IDH1 (H315 IDH1), and mutation of this residue to alanine destroys catalytic activity because this residue binds the phosphate of NADP^+^ (12,64,65). Low pH also increases rates of NADPH consumption in oxen IDH2, and it is suggested that this is driven, at least in part, by better CO_2_ solubility at lower pH driving the reverse reaction (66). Thus, some reactions catalyzed by IDH1 in various organisms appear to be sensitive to pH under select conditions, but a lack of reported steady-state parameters and inconsistent reaction conditions complicate conclusions.

Here we present a thorough analysis of IDH1 catalysis at varying pH values and show that the forward reaction appears to be regulated by pH. The *k*_cat_ of the forward reaction (isocitrate to αKG) shows the most reliable pH dependence in the buffer systems tested here, though the pH-sensitivity of kinetic parameters of the reverse reaction is buffer-dependent. We identified buried networks of acidic and basic residues that included D273 and K217 IDH1, two residues that were calculated to have shifted pK_a_ values into the physiological range. When D273 IDH1 was mutated to non-ionizable residues, there was a drastic loss of activity and decreased sensitivity to pH and, in the R132H IDH1 background, apparent disruption of mutant IDH1 inhibitor binding. Overall, this work uses structural informatics and experimental methods to identify and evaluate a mechanism of pH-dependent catalytic regulation that is mediated, at least in part, by a buried acidic residue found in the IDH1 dimer interface.

## RESULTS

### Effects of pH modulation on WT IDH1 activity

Heterologously expressed and purified human IDH1 was characterized under steady-state parameters to determine the effects of pH on catalysis. The *k*_cat_ of the forward reaction, isocitrate to αKG, in both KPhos and tris/bis-tris buffers was pH dependent, exhibiting a trend that increased with increasing pH. However, variability in *K*_m_ minimized this trend when comparing catalytic efficiencies (Fig. 2, Table 1), suggesting substrate titration may also affect catalysis.

**Table 1.** Kinetic parameters for the normal forward and reverse reactions catalyzed by WT IDH1 in varying pH and buffers. Steady-state rates were derived from fitting plots of *k*_obs_ versus substrate concentration with the Michaelis-Menten equation, with the standard error of the mean (S.E.M.) determined from the deviance from these hyperbolic fits. Data were obtained as described in Fig 2.

| IDH1 | Buffer | Reaction | $k_{\text{cat}}$, s<sup>-1</sup> | $K_{\text{m}}$, mM | $k_{\text{cat}}/K_{\text{m}}$, mM<sup>-1</sup>s<sup>-1</sup> |
| --- | --- | --- | --- | --- | --- |
| WT | Bis-tris, pH 6.2 | Isocitrate → αKG | 20.0 ± 0.4 | 0.042 ± 0.004 | (0.48 ± 0.05) × 10<sup>3</sup> |
| WT | Bis-tris, pH 6.5 | Isocitrate → αKG | 23.1 ± 0.6 | 0.048 ± 0.006 | (0.48 ± 0.06) × 10<sup>3</sup> |
| WT | Bis-tris, pH 7.0 | Isocitrate → αKG | 32 ± 1 | 0.075 ± 0.009 | (0.43 ± 0.05) × 10<sup>3</sup> |
| WT | Tris, pH 7.2 | Isocitrate → αKG | 39 ± 1 | 0.11 ± 0.01 | (0.35 ± 0.03) × 10<sup>3</sup> |
| WT | Tris, pH 7.5 | Isocitrate → αKG | 40.4 ± 0.8 | 0.030 ± 0.003 | (1.4 ± 0.1) × 10<sup>3</sup> |
| WT | Tris, pH 7.8 | Isocitrate → αKG | 35.4 ± 0.6 | 0.035 ± 0.003 | (1.01 ± 0.09) × 10<sup>3</sup> |
| WT | Tris, pH 8.0 | Isocitrate → αKG | 38.3 ± 0.9 | 0.038 ± 0.004 | (1.0 ± 0.1) × 10<sup>3</sup> |
| WT | KPhos, pH 6.2 | Isocitrate → αKG | 20.2 ± 0.4 | 0.027 ± 0.003 | (0.75 ± 0.08) × 10<sup>3</sup> |
| WT | KPhos, pH 6.5 | Isocitrate → αKG | 23.0 ± 0.4 | 0.034 ± 0.003 | (0.68 ± 0.06) × 10<sup>3</sup> |
| WT | KPhos, pH 6.8 | Isocitrate → αKG | 27.5 ± 0.6 | 0.040 ± 0.004 | (0.69 ± 0.07) × 10<sup>3</sup> |
| WT | KPhos, pH 7.0 | Isocitrate → αKG | 31.7 ± 0.8 | 0.050 ± 0.006 | (0.63 ± 0.08) × 10<sup>3</sup> |
| WT | KPhos, pH 7.2 | Isocitrate → αKG | 35.5 ± 0.8 | 0.060 ± 0.006 | (0.59 ± 0.06) × 10<sup>3</sup> |
| WT | KPhos, pH 7.5 | Isocitrate → αKG | 37.6 ± 0.7 | 0.061 ± 0.005 | (0.62 ± 0.05) × 10<sup>3</sup> |
| WT | KPhos, pH 7.8 | Isocitrate → αKG | 40.0 ± 0.7 | 0.066 ± 0.005 | (0.61 ± 0.05) × 10<sup>3</sup> |
| WT | KPhos, pH 8.0 | Isocitrate → αKG | 39.2 ± 0.7 | 0.067 ± 0.005 | (0.59 ± 0.05) × 10<sup>3</sup> |
| WT | Bis-tris, pH 6.2 | αKG → isocitrate | 1.6 ± 0.1 | 0.04 ± 0.01 | 40 ± 10 |
| WT | Bis-tris, pH 6.5 | αKG → isocitrate | 3.2 ± 0.2 | 0.12 ± 0.04 | 27 ± 9 |
| WT | Bis-tris, pH 6.8 | αKG → isocitrate | 3.3 ± 0.2 | 0.14 ± 0.03 | 24 ± 5 |
| WT | Bis-tris, pH 7.0 | αKG → isocitrate | 3.8 ± 0.3 | 0.14 ± 0.04 | 27 ± 8 |
| WT | Tris, pH 7.2 | αKG → isocitrate | 3.4 ± 0.3 | 0.26 ± 0.07 | 13 ± 4 |
| WT | Tris, pH 7.5 | αKG → isocitrate | 3.6 ± 0.2 | 0.246 ± 0.06 | 15 ± 4 |
| WT | KPhos, pH 6.2 | αKG → isocitrate | 3.5 ± 0.3 | 0.6 ± 0.2 | 6 ± 2 |
| WT | KPhos, pH 6.5 | αKG → isocitrate | 4.1 ± 0.2 | 1.0 ± 0.1 | 4.1 ± 0.5 |
| WT | KPhos, pH 6.8 | αKG → isocitrate | 4.9 ± 0.2 | 0.65 ± 0.08 | 7.5 ± 0.1 |
| WT | KPhos, pH 7.0 | αKG → isocitrate | 4.1 ± 0.2 | 0.71 ± 0.09 | 5.8 ± 0.8 |
| WT | KPhos, pH 7.2 | αKG → isocitrate | 3.87 ± 0.07 | 0.28 ± 0.02 | 14 ± 1 |
| WT | KPhos, pH 7.5 | αKG → isocitrate | 3.8 ± 0.1 | 0.46 ± 0.05 | 8.3 ± 0.9 |
| WT | KPhos, pH 7.8 | αKG → isocitrate | 1.87 ± 0.08 | 0.87 ± 0.09 | 2.1 ± 0.2 |

**Figure 2.**
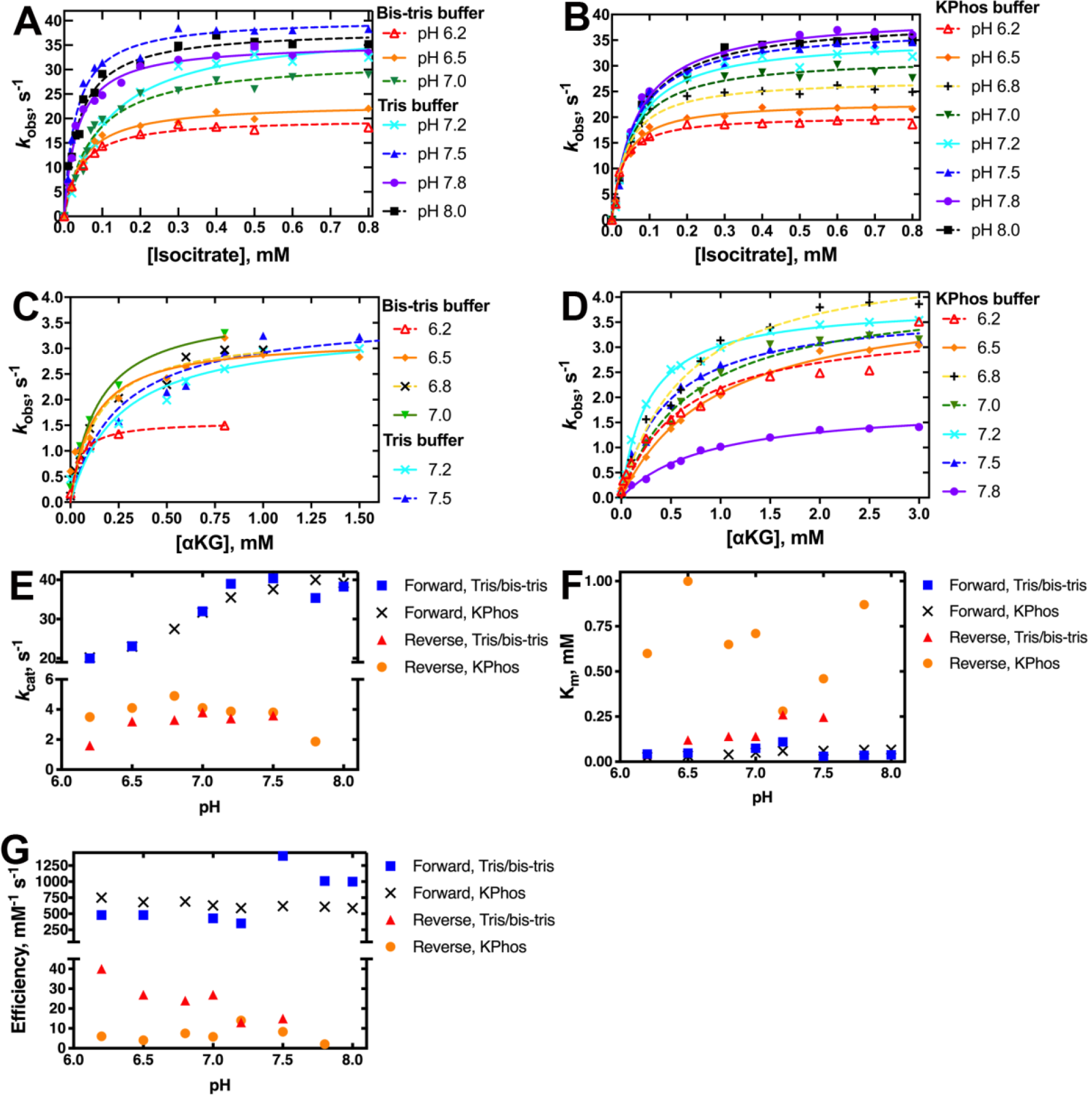
Effects of pH and buffer on WT IDH1 catalysis. Steady-state rates of the forward reaction, the conversion of isocitrate to αKG, catalyzed by WT IDH1 were measured as function of varying pH using either tris (pH 7.2-8.0) or bis-tris (pH 6.2-7.0) buffers (A), or KPhos buffer (B). Steady-state rates of the reverse reaction, the conversion of αKG to isocitrate, catalyzed by WT IDH1 were measured as a function of varying pH using either tris or bis-tris buffers (C), or KPhos buffer (D). In (A-D), at least two protein preparations were used to measure the observed rate constants (*k*_*obs*_) at varying substrate concentrations, which were calculated from determining the linear portion of plots of concentration versus time. Each point in the curve represents a single replicate. The variation in *k*_cat_ (E), *K*_m_ (F), and *k*_cat_/*K*_m_ (efficiency) (G) due to change in pH from parts (A-D) is shown.

The reverse reaction catalyzed by human WT IDH1 is generally much less efficient than the forward reaction, resulting primarily from a decrease in *k*_cat_ but also from an increase in *K*_m_ particularly in KPhos buffer (Fig. 2C-G). Here, any pH dependence observed was primarily limited to studies in tris/bis-tris buffers, as only a very slight trend was seen in *k*_cat_ in KPhos buffer. Modest decreases in *K*_m_ at more acidic pH values were observed primarily in tris/bis-tris buffers, and limited effects on *k*_cat_ mean this small trend was also seen when comparing catalytic efficiencies (Fig. 2E-G, Table 1). pH-dependent trends for the reverse reaction were relatively small and show much greater buffer-to-buffer variability.

To ensure that changes in activity were not due to protein unfolding, we used circular dichroism (CD) to show that IDH1 secondary structure features remained stable through this range of pH values, with no significant change in *T*_m_ value (Fig. S1). Thus, from our steady-state assessment, we concluded that the pH dependence of *k*_cat_ for the forward reaction had the most consistent trend in our assay conditions compared to the reverse and neomorphic reactions, which had more modest trends and were typically buffer-dependent. Consequently, the forward reaction became the major focus of our consequent characterization.

### Modulation of pH in cell lines

Small but meaningful changes in intracellular pH can occur during a variety of cellular processes like apoptosis (26), under changing environments like hypoxia (67), and in disease states like neurodegenerative diseases (34,35), cancer (25,36-39), and diabetes (68). Since an increase in pH led to increased *k*_cat_ values for the forward reaction in biochemical assays with WT IDH1, we sought to determine if acidic cellular environments led to changes in IDH1-relevant metabolite levels.

We modulated the cellular pH_i_ by treating cells with proton transport inhibitors or ion exchange inhibitors (69,70) and then quantified metabolite levels in cells using gas chromatography coupled to mass spectrometry (GC/MS). These experiments were admittedly limiting since several metabolic pathways affect isocitrate and αKG levels. In an attempt to mitigate this, we selected isogenic cell lines that had varying levels of WT IDH1: patient-derived HT1080 cells containing either an endogenous heterozygous R132C mutation (R132C/+ IDH1, where + denotes WT IDH1) or with an R132C-ablated version of this cell line that also stably overexpresses WT IDH1 (-/+++ IDH1) (71). As a result of the R132C IDH1 mutation, D2HG is produced and WT IDH1 activity is ablated in protein expressed from this allele (22,72,73). A series of proton transport inhibitors, including esomeprazole (ESOM), which targets proton pumps like V-ATPase, and ion exchange inhibitors 4,4’-diisothiocyano-2,2’-stilbenedisulfonic acid (DIDS) and 5-(*N*-ethyl-*N*-isopropyl)amiloride (EIPA), were tested at varying concentrations to find the most effective methods to decrease pH_i_ without causing an observed increase in cell death. For the HT1080 cell lines, treatment with 200 µM ESOM was most effective (Table 2, Fig. 3A).

**Table 2.**
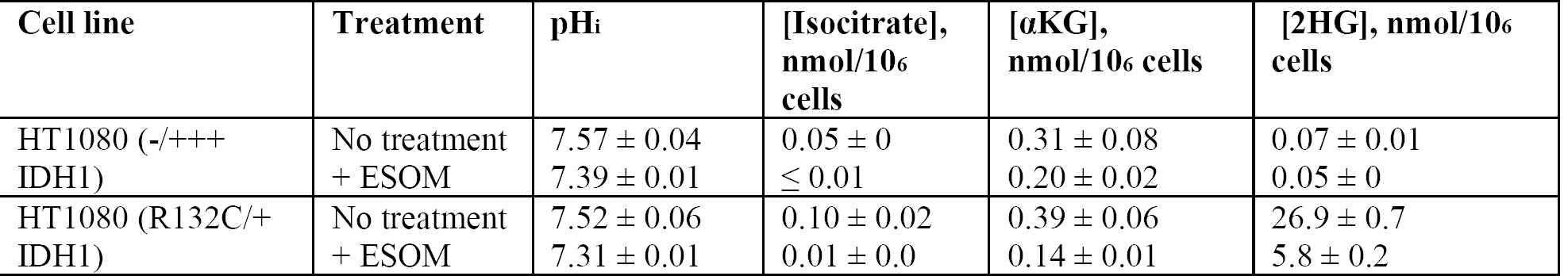
Cellular quantitation of metabolites related to IDH1 activity at varying pH, biological replicates of two.

**Figure 3.**
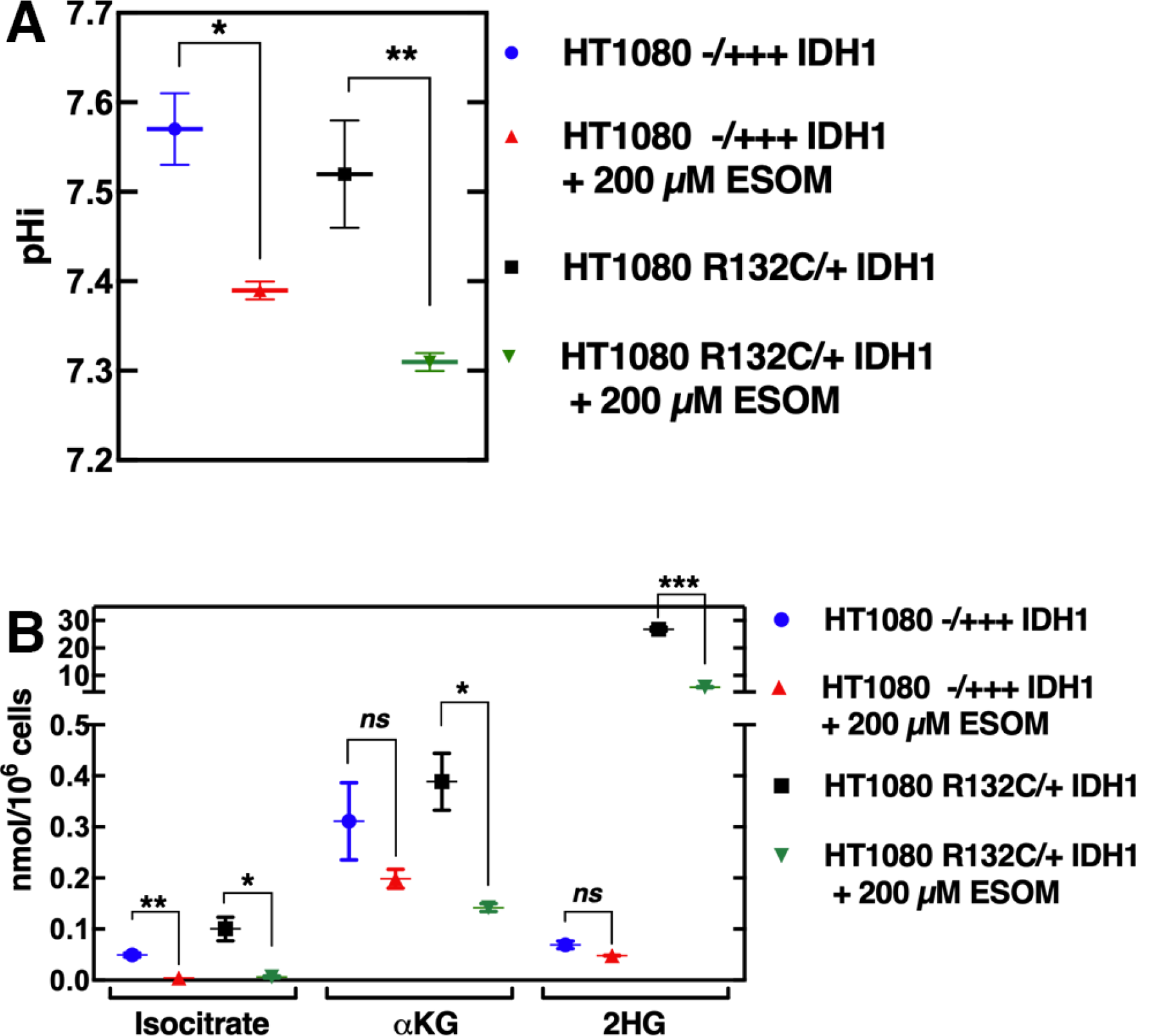
Change in IDH1-related metabolites upon pH_i_ modification. Two sets of cell lines were used: patient-derived HT1080 cells, which contain an endogenous heterozygous R132C IDH1 mutation (R132C/+ IDH1), or HT1080 cells where the R132C IDH1 allele was ablated with a T77A mutation to destroy neomorphic activity (αKG to D2HG production) in the R132C allele, and made to stably overexpress WT IDH1 (-/+++ IDH1) (71). Cell lines were either untreated or treated with 200 µM ESOM to decrease the pH_i_. The resulting change in pH are shown in (A). Experiments were performed as technical triplicates. (B) Metabolites were then quantified and changes in isocitrate, αKG, and 2HG (isomers unresolved) are highlighted (see Table S1 for additional metabolites). Experiments were performed as biological duplicates. *Not significant (ns)* > 0.05, \**p* ≤ 0.05, \*\**p* ≤ 0.01, \*\*\**p* ≤ 0.001.

Isocitrate levels significantly dropped upon ESOM treatment in HT1080 cells (Table 2, Fig. 3B), though these values were near the limit of detection. αKG also decreased upon a shift to an acidic pH_i_ in HT1080 cell lines (Table 2, Fig. 3B), though significance wasn’t achieved in the case of the HT1080 -/+++ IDH1 cells. A decrease in αKG concentration was not surprising due to the observed decrease in *k*_cat_ in more acidic buffers, though of course many enzymes contribute to αKG levels, including IDH2 and IDH3. Notably, a global decrease in metabolite concentrations was not observed; fifty common metabolites were quantified in each of the cell lines, and ESOM treatment yielded both increases and decreases in metabolite levels (Table S1), minimizing the possibility that cells were simply in the process of dying. 2HG levels in the mutant cell line (the *D* and *L* isomers of 2HG cannot be resolved in these experiments), though not a focus in this work, were also noted to decrease upon ESOM treatment.

### Characterizing IDH1 ionizable networks

To explore possible mechanisms for the pH dependence observed in the forward reaction catalyzed by IDH1, we used a structural informatics approach known as pHinder (29,43,74) to identify potential pH-sensing ionizable residues. Briefly, the pHinder algorithm uses triangulation to calculate topological networks of ionizable residues (D, E, H, C, K, and R) in proteins. These networks identify characteristics that can be predictive of protein structure-function relationships. For example, portions of ionizable networks buried inside proteins or that comprise contiguous stretches of acidic or basic residues tend to contain residues with pK_a_ values shifted to the physiological pH range (pH 5 to 8). Small changes in pH may then be sensed by these residues via a change in their ionization state, resulting in structural and thus functional changes (28,29,51,75).

We identified a striking number of buried acidic and basic residue networks in structures of WT IDH1 complexed to NADP^+^ (apo, 1T09 (76)), WT IDH1 bound to NADP^+^, isocitrate, and Ca^2+^ (holo, 1T0L (76)), and R132H IDH1 bound to αKG, NADPH, and Ca^2+^ (holo, 4KZO (64)) (Figs. 4, S2). Here, Ca^2+^ mimics the catalytic Mg^2+^ metal required for catalysis. We will use the nomenclature of X###, where X refers to the single amino acid code for the WT residue present at residue number ###. As IDH1 is a dimer, we will distinguish the two polypeptide chains by using X### and X###’. Mutations will be noted as X###Y, where X at position ### becomes mutated to Y.

**Figure 4.**
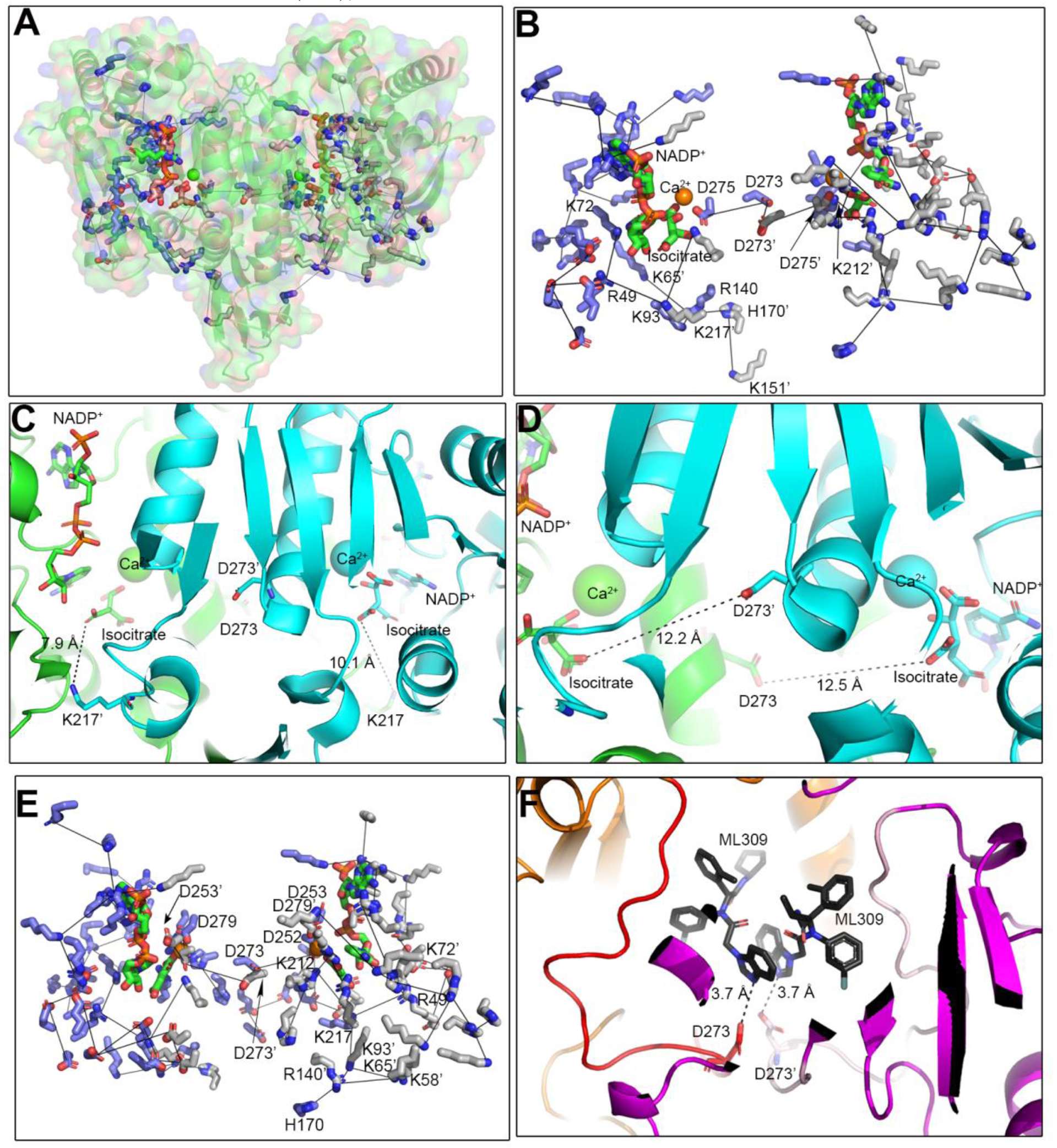
pHinder analysis of the holo IDH1 dimer. (A) Networks of buried ionizable residues were identified using pHinder in a previously solved structure of holo WT IDH1 bound to NADP_+_, isocitrate, and Ca_2+_ (1T0L (76)). These networks are shown as grey lines and with the residues making up this network shown as purple (chain A) or grey (chain B) sticks. (B) Only the residues involved in this network, as well as isocitrate and NADP_+_ (shown in green) are highlighted. (C) The location of K217 (chain A) and K217’ (chain B) relative to the active sites. (D) The location of D273 (chain A) and D273’ (chain B) relative to the active sites. (E) pHinder was also used to identify a network of buried ionizable residues in a previously solved structure of holo R132H IDH1 bound to αKG, NADP_+_, and Ca_2+_ (4KZO (64)). For clarity, only residues described in the text as being part of networks involving K212 and D273 are labeled in part B and (F) Two ML309 ligands (72) were docked into the density present in a cryo-EM structure of R132C IDH1 incubated with ML309 (115), and the localization of D273 relative to these inhibitors is shown.

Of note, in the holo form of WT IDH1, a short acidic network involving buried residues D275, D273, D273’, and D275’ IDH1 traversed through the middle of the IDH1 dimer (Figs. 4B, S2). Residues D275 and D275’ in IDH1 are involved in ligand and metal binding, but D273 and D273’ are too distant for any direct interactions with substrates (Fig. 4D). In the apo form of WT IDH1 (1T09 (76)), D273 and D273’ were the only amino acids making up a very short network chain, though interestingly R132 in one chain localized to a similar position as D273 in the holo structure (Fig. S2B).

Given the unusual number and complexity of the buried ionizable networks in IDH1, we chose to focus on two networks, a short acidic network that traversed through the middle of the IDH1 dimer, and a longer basic network within each IDH1 monomer (Figs. 4, S2). The core component of the short acidic network involves residues D273 and D273’. These residues are found at the dimer interface of IDH1, far from the enzyme active site and close to where allosteric mutant IDH1 inhibitors bind (Fig. 4F). The D273 residues also participated in a longer network involving D253, D273’, D279, D273, D252, D252’, D253’, and D279’ in the structure of the holo R132H IDH1 (4KZO (64)) (Fig. 4E), though D275 was no longer involved.

In contrast to the short acidic network, we also identified a longer basic network consisting of K151’, H170’, R140, K65, K93, K217’, K212’, R49, and K72 IDH1 in holo WT IDH1 (1T0L (76)), Fig. 4B). This long basic network intersected with the active site, as K212’ is a catalytic residue. Some side chains in the network were buried deeply while other residues were partially exposed. To avoid direct mutational effects on IDH1 catalytic activity (i.e., affecting directly affecting substrate binding or chemistry), we focused our attention on one of the most deeply buried side chains, K217’, that was more distant from the active site (Fig. 4C). The A-chain K217 residue also appeared in an additional network containing 11 amino acids (R100’, H170, K217, R132’, K93’, R109’, R140’, R49’, K58’, K72’, and K212 IDH1), which involved most of the same residues in the opposite chains except R100, R109, R132 (all metal/substrate coordinating residues), and K58; instead K151’ and K65 were included (Fig. 4B).

Having selected D273 and K217 residues for in-depth analysis, we next calculated the pK_a_ value of each residue in the WT IDH1 structure (1T0L (76)) using PROPKA (77,78). The objective of these calculations was to provide a rough estimate of the D273 and K217 pK_a_ values (Table S2). K217 had a calculated pK_a_ value of 7.97, while K217’ had a calculated pK_a_ value of 8.24. This indicated a downshifts towards more acidic, physiologically-relevant pK_a_ values, consistent with previous findings that buried K residues can have downshifted pK_a_ values that can tune protein stability (40,42). D273 and D273’ were also predicted to have more physiologically relevant pK_a_ values, with upshifts to 6.59 and 6.4, respectively.

Based on our observations using structural informatics and pK_a_ calculations, we hypothesized that D273 and K217 in IDH1 had features consistent with pH-sensing residues, namely buried ionizable side chains with pK_a_ values shifted to more physiological ranges. Of note, unlike K, D, and E residues, R residues have not been observed to have pK_a_ value shifts when buried (79). Indeed, we show minimal changes in pK_a_ in R residues (Table S2). As a result of these cumulative findings, we sought to confirm the role of D273 and K217 in IDH1 catalysis in both biochemical and cell-based experiments.

### Residue K217 in IDH1 has only a modest role in catalysis

We first ablated the ability of the D273 and K217 residues to undergo changes in ionization state. K217 IDH1 is found in a loop located between the β10 β-sheet and α7 α-helix, ∼8-10 Å from isocitrate depending on the chain (Fig. 4C). As this residue in both monomer chains was predicted to have a downshifted pK_a_ value from ∼10.7 in solution to 7.97 and 8.24, we selected mutations that would introduce minimal structural/steric changes but ablate ionizability. Methionine, a nonpolar residue, and glutamine, a polar uncharged residue, were selected. Recombinant K217M and K217Q IDH1 were heterologously expressed and purified from *E. coli*, and we measured the catalytic efficiency of the conversion of isocitrate to αKG in steady-state kinetic conditions. K217M IDH1 had no effect on *k*_cat_ and a modest increase in *K*_m_, yielding only a 2.2-fold decrease in catalytic efficiency (Fig. 5A, Table 3). K217Q was more disruptive, with a 5.4-fold decrease in catalytic efficiency driven primarily through a 4-fold increase in *K*_m_ (Fig. 5A, Table 3). Overall, ionizability at this residue did not appear vital for catalysis, though its mutation did affect activity.

**Table 3.** Kinetic parameters for the normal forward and neomorphic reactions catalyzed by IDH1 variants. Steady-state rates of *k*_cat_, *K*_m_, and *k*_cat_/*K*_m_ were derived from fitting plots of *k*_obs_ versus substrate concentration with the Michaelis-Menten equation, with the standard error determined from the deviance from these hyperbolic fits. Data were obtained as described in Fig 5.

| IDH1, reaction | $k_{\text{cat}}$, s<sup>-1</sup> | $K_{\text{m}}$, mM | $k_{\text{cat}}/K_{\text{m}}$, mM<sup>-1</sup>s<sup>-1</sup> |
| --- | --- | --- | --- |
| WT, isocitrate → αKG | 37.6 ± 0.7 | 0.061 ± 0.005 | 620 ± 50 |
| K217M, isocitrate → αKG | 42 ± 1 | 0.15 ± 0.02 | 280 ± 40 |
| K217Q, isocitrate → αKG | 28.7 ± 0.5 | 0.25 ± 0.02 | 115 ± 9 |
| D273N, isocitrate → αKG | 22 ± 1 | 19 ± 2 | 1.2 ± 0.1 |
| D273S, isocitrate → αKG | 15 ± 1 | 12 ± 3 | 1.3 ± 0.3 |
| D273L, isocitrate → αKG | 7.0 ± 0.2 | 1.9 ± 0.2 | 3.7 ± 0.4 |
| R132H, αKG → D2HG | 1.44 ± 0.05<sup>a</sup> | 1.5 ± 0.2<sup>a</sup> | 1.0 ± 0.1<sup>a</sup> |
| D273N/R132H, αKG → D2HG | ≤ 0.02<sup>b</sup> | N.D.<sup>c</sup> | N.D.<sup>c</sup> |
| D273L/R132H, αKG → D2HG | ≤ 0.02<sup>b</sup> | N.D.<sup>c</sup> | N.D.<sup>c</sup> |
| D273S/R132H, αKG → D2HG | ≤ 0.02<sup>b</sup> | N.D.<sup>c</sup> | N.D.<sup>c</sup> |

<sup>a</sup> From Ref. (116).
<sup>b</sup> No detectable activity (NADPH consumption) was observed, so a limit of detection is listed. Here, 500 nM of protein and 1-5 mM αKG was used.
<sup>c</sup> Due to no detectable activity, $K_{\text{m}}$ and $k_{\text{cat}}/K_{\text{m}}$ values could not be determined (N.D., not determined).

**Figure 5.**
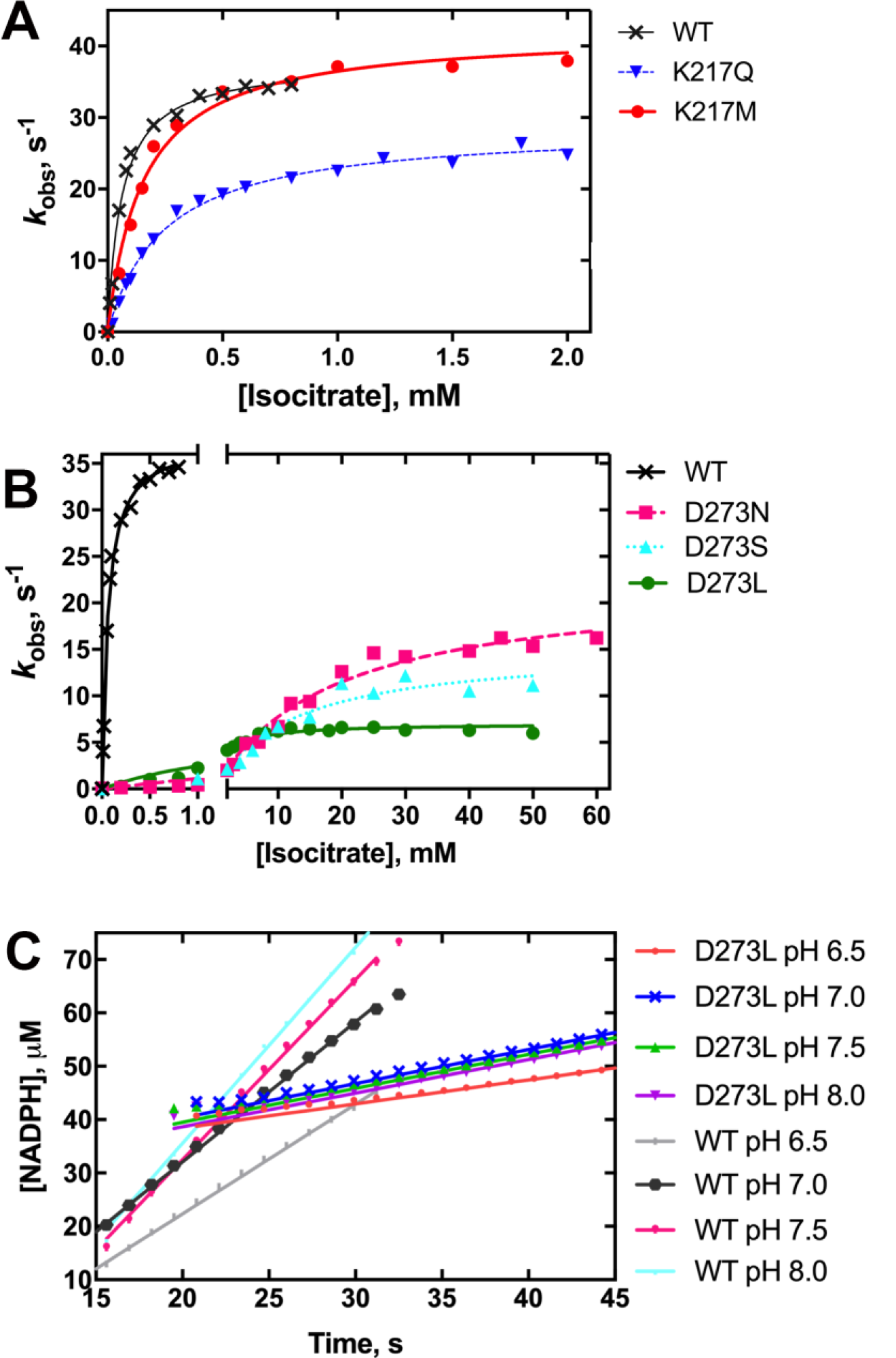
Kinetic characterization of K217 and D273 mutants in the WT IDH1 background. Steady-state rates of the normal forward reaction, the conversion of isocitrate to αKG, as catalyzed by (A) K217Q and K217M IDH1, and (B) D273N, D273S, and D273L IDH1 are shown relative to WT IDH1. *k*_obs_ values were obtained as described in Fig. 2. Replicates were performed for higher concentrations of isocitrate in the case of D273S IDH1 as these measurements proved more error prone. (C) Comparison of sensitivity to pH of the normal forward reaction in D273L and WT IDH1. Four representative pH values were tested, and loss of pH sensitivity was observed in D273L for most of the pH ranges tested. The average slope for the D273L mutants is 0.00042 ± 0.00006 µM s_-1_, while the average slope is 0.04 ± 0.01 µM s_-1_, indicating higher variability for the latter enzyme, and loss of pH dependence for the former. Each point in the curve represents a single replicate, and at least two different protein preparations were used to obtain replicates in the curves.

### Residue D273 in IDH1 appears to play a role in pH-regulated catalysis and inhibitor binding

D273 in IDH1 was predicted to have an upshifted pK_a_ from a standard value of ∼3.7 in solution to a more physiologically relevant value of ∼6.5. This residue is located in the first third of the α10 helix, an important regulatory domain that transitions from an ordered loop in the apo form of IDH1 to a helix in the holo form (76). This residue is ∼12 Å from the nearest substrate (isocitrate) and is found at the dimer interface (Fig. 4D). Again, mutations were selected that would minimally disrupt the overall structure but would destroy ionizability, and thus D273N, D273L, and D273S IDH1 were generated. After heterologously expressing and purifying these mutants, we found that the kinetic profile of αKG production (the normal forward reaction) was severely affected (Fig 5B, Table 3). Insertion of a nonpolar residue at this position had very drastic effects, with D273L IDH1 exhibiting a ∼170-fold decrease in catalytic efficiency, driven by a 5.4-fold decrease in *k*_cat_ and 31-fold increase in *K*_m_. Interestingly, the more conservative D273N and D273S IDH1 mutations had even more deleterious effects. D273N IDH1 had a >500-fold decrease in *k*_cat_/*K*_m_, driven primarily through >300-fold increase in *K*_m_. D273S IDH1 had a similar effect on *k*_cat_ and *K*_m_ as D273N IDH1 (∼2.5-fold and nearly 200-fold decreases, respectively, relative to WT IDH1). Thus, D273 IDH1 mutations produced very drastic decreases in catalytic efficiency for the forward reaction as compared to WT IDH1.

To determine whether mutation of the D273 residue in IDH1 affected the observed pH sensitivity, we measured the rates of isocitrate to αKG conversion by D273L IDH1 at saturating concentrations of substrates in KPhos buffer at pH 6.5, 7.0, 7.5, and 8.0 (Fig. 5C). Incubating WT IDH1 with 600 µM isocitrate and 200 µM NADP^+^ yielded *k*_obs_ of 23.0, 27.5, 37.6 and 39.2 s^-1^ respectively, showing clear trends in pH dependency. In contrast, D273L IDH1 catalysis was not altered by changes in pH (i.e. was insensitive to pH), except at the most acidic environment; *k*_obs_ rates of 4.5, 6.4, 6.4, and 6.3 s^-1^ at pH 6.5, 7.0, 7.5, and 8.0, respectively, were observed (Fig. 5C), supporting our hypothesis that D273 is a critical mediator contributing to pH sensitivity.

The α10 regulatory domain has been posited to be important for conferring selectivity for mutant IDH1 inhibitor binding (80). As D273 is located in this domain, we next asked if D273 mutants altered catalysis and inhibitor binding in the R132H IDH1 background. Double mutants (D273N/R132H, D273S/R132H, and D273L/R132H IDH1) were expressed and purified, and the steady-state rates of conversion of αKG to D2HG were measured. However, these double mutants were essentially catalytically inactive. Thus, we only report a limit of detection, *k*_obs_, of ≤0.02 s^-1^, as measured rates fell below this value. Rates were below the detectable limit despite sampling a range of enzyme concentrations (up to 500 nM) across 4 different recombinant protein preparations with high substrate concentration (up to 5 mM αKG) (Table 3) to improve detection of product formation. This low activity precluded our determination of *K*_m_, *k*_cat_/*K*_m_, or IC_50_ measurements. Instead, we used isothermal titration calorimetry (ITC) to determine the affinity of D273L/R132H IDH1 for two commercially available selective mutant IDH1 inhibitors, AGI-5198 (81) and ML309 (72). Interestingly, both inhibitors showed that binding was affected. We measured a *K*_d_ of 3.3 +/- 0.5 µM for AGI-5198 binding to IDH1 D273L/R132H IDH1 (compared previously reported IC_50_ value of 0.07 µM for R132H IDH1 (81)). Under the conditions of our experiments, no binding of ML309 to D273L/R132H IDH1 was detected, which implied no binding or binding with a *K*_d_ <20 µM due to limits of detection (versus previously reported *K*_d_ value of 0.48 ± 0.05 µM for R132H IDH1(82)) (Table 3). In sum, this indicates that D273 plays in important role in WT and mutant IDH1 catalysis, pH-sensing in catalysis of the forward normal reaction, and in mutant IDH1 inhibitor binding.

## DISCUSSION

Here we report an assessment of the effects of pH on IDH1 catalysis. We show that WT IDH1 is sensitive to pH in catalyzing the forward reaction (Fig. 2), though increases in *k*_cat_ as the pH increases are mitigated by some corresponding increases in *K*_m_, leading to essentially no pH-dependence when considering catalytic efficiency (*k*_cat_/*K*_m_, Table 1). As mammalian cellular concentrations of isocitrate are near the *K*_m_ values we report, it is possible that local concentrations of isocitrate would support rates near *k*_cat_ (83), making this a physiologically relevant finding.

Only the reverse reaction performed in tris/bis-tris buffer showed pH-dependence in catalytic efficiency (Table 1). The buffer-dependence of these findings suggests that previous reliance on a decrease in pH to drive the reverse normal reaction (αKG to isocitrate) by us and others (62) is an effective *in vitro* strategy for studying catalytic features of IDH1. However, it is likely not appropriate to extrapolate that local changes in cellular pH affect rates of the reverse reaction as physiological cellular changes in pH may be too narrow. It has previously been shown that hypoxia is effective at driving lipid biosynthesis via isocitrate production by IDH1 (18). In addition to low oxygen content, hypoxia is associated with a decrease in local pH_i_ (for example, (84-86)). However, based on our data, we predict factors other than pH-regulated catalysis are driving switching to reductive metabolism. This may include changes in substrate levels, regulation of IDH1 function as a result of hypoxia, etc.

There have been other reported cases of metabolic enzymes having buffer-dependent activities. For example, it has been posited that inorganic phosphate can activate fumarase at higher concentrations, and inhibit this enzyme at concentrations <5 mM (87,88). Our experiments did not test whether phosphate acts as an allosteric activator in IDH1, though this will be an interesting future direction to explore. Certainly, generating CO_2_ as a substrate as required to study the reverse reaction can affect pH if buffering capacity is not sufficient. Further, and perhaps more importantly, CO_2_ may react with the amines of tris and bis-tris buffers (89), affecting the local concentration of CO_2_ available to drive the reverse normal reaction. Of note, our buffer testing was limited to KPhos and tris-based buffers, examples of inorganic and non-inorganic ionic buffers. We did not include zwitterionic buffers like HEPES or MOPS, which can have superior buffering capacity in cell-based studies. However, unlike KPhos and Tris buffers, HEPES and MOPS are more redox-sensitive, and thus can introduce new challenges when studying redox-sensitive reactions. However, future work should include these buffering systems.

In addition to the acid and base chemistry required for IDH1 catalysis (90,91), the ionization state of the substrates themselves are important for binding. This can also be playing a role in the results reported here. In IDH purified from pig heart (isoform not specified), isocitrate preferably binds when isocitrate has all three carboxyl groups ionized (deprotonated) (92-95). It will be interesting to explore in future work if the wide variation in *K*_m_ values we observe (Fig. 2) is related to this issue of altered ionic form of the substrates.

The use of steady-state kinetics in this work to investigate pH sensitivity means that we are examining the role of pH on the overall rate-limiting step (predicted to be hydride transfer (90,91,96,97)), but are potentially missing any sensitivity to pH in non-rate-limiting steps. In order to probe the role of pH in additional steps of catalysis, pre-steady-state kinetic methods would be required, which would be an interesting future direction. There is precedence for pH sensitivity in NADP(H)-dependent hydride transfer reactions; for example, *E. coli* dihydrofolate reductase (DHFR) is an interesting case where the rate-limiting step changes from product release at low pH to hydride transfer at high pH (98), with hydrogen bonds in key loop conformations driving much of these observations (99-102).

Using the structural informatics program pHinder, we identified the buried ionizable residues K217 and D273 in IDH1 as candidates for having pH-sensing properties. Experimentally, we found that the ionizability of D273 affected both catalysis and pH sensitivity. This was not the case for K217, which we speculate is likely more important for tuning protein stability as described previously for buried K residues in other proteins (40,42). These findings demonstrate the advantage of combining structural informatics with experimental validation. Importantly, these findings also confirm the consistent utility of the pHinder algorithm for identifying ionizable residues that are unexpectedly important for both pH sensing and protein function, as D273 is quite distant from the active site. As a result, its importance in IDH1 function would be difficult to predict by other means. Indeed, in this case, we identified a possible role for D273 in pH-sensitive catalysis, supported by our data showing loss of pH-dependent catalysis for the D237L IDH1 mutant in physiologically relevant ranges. As a drop in *k*_cat_ was only observed at pH 6.5 for D273L, catalysis at even lower pH values could be tested, though low pH values lose physiological relevance and may adversely (and artifactually) affect protein stability. More importantly, we feel exploring additional D273 mutants such as D273E to identify the requirement for a titratable carboxylate residue at this position as a requirement for pH-sensing would be worthy of future exploration.

D273 is conserved in IDH1 and IDH2 (D312 IDH2) and is found in the α10 regulatory domain (using IDH1 nomenclature) (76). According to simple modeling mutational variants D273N, D173S, and D273N are not predicted to drastically affect either the global structure of IDH1 (Fig. 6A) or even local folding features (Fig. 6B, 6C). The regulatory role of this domain likely differs when comparing WT and mutant IDH1, where in the latter form of the protein, unraveling of the α10 helix (80) and changes in water dynamics (82) have been posited to contribute to selectivity of inhibitor binding to mutant over WT IDH1. In contrast, in structures of mutant IDH2, this domain remains in the helical conformation in apo, holo, and inhibitor-bound forms (80,103). Interestingly, the D273 homolog in IDH2, D312, is located on the same side of the helix and only 5.3 Å from Q316 IDH2, a residue involved in acquired resistance to mutant IDH2 inhibitor binding. Q316E IDH1 has been identified in patients treated with the selective mutant IDH2 inhibitor Enasidenib (103), and results in resistance to this drug (104). Thus, our reported decrease in affinity for mutant IDH1 inhibitors is not surprising when considering the proximity of D273 to the allosteric pocket (Fig. 6D).

**Figure 6.**
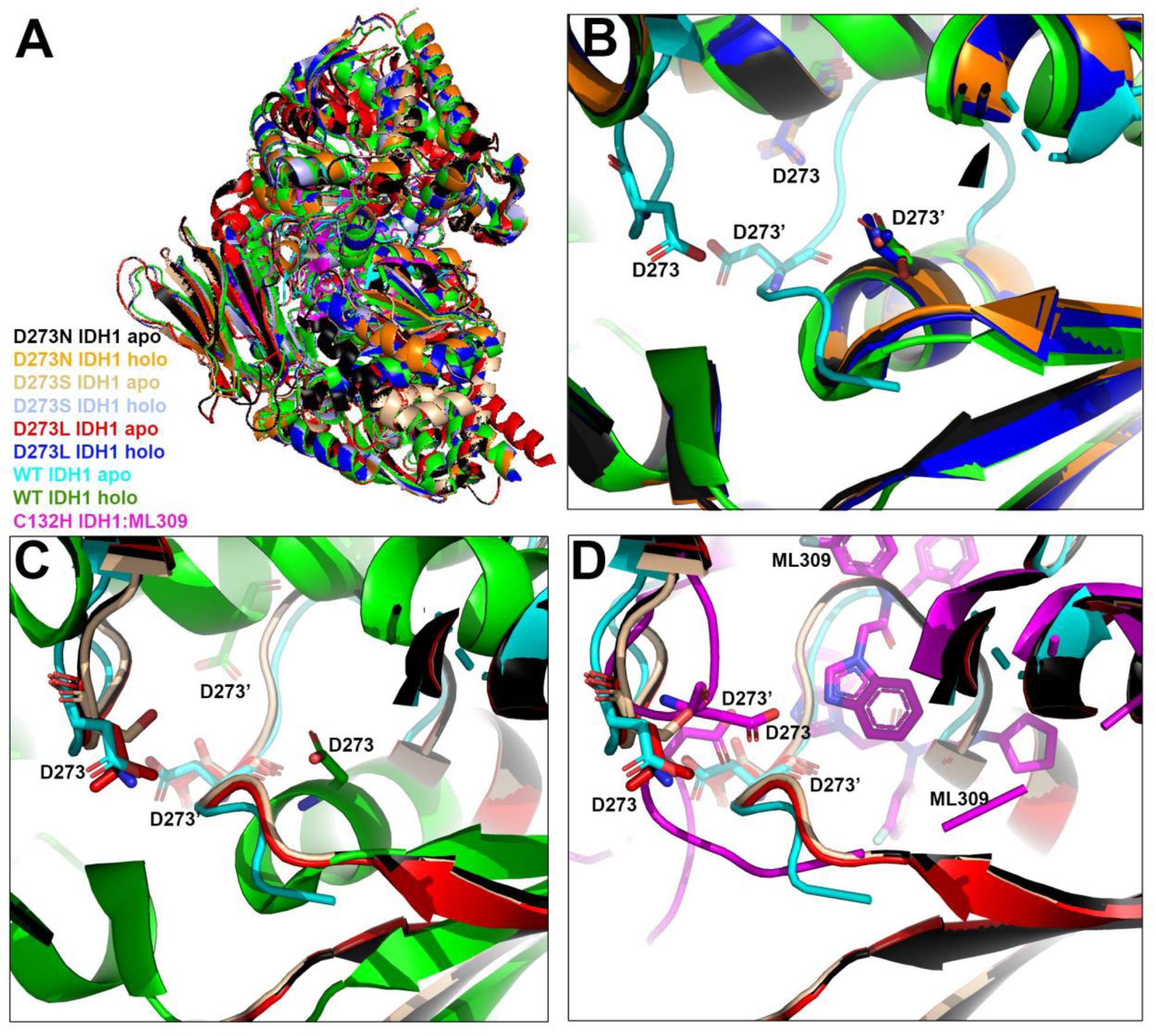
Structural models of the D273 mutant series. (A) The orientation of IDH1 as shown in parts B-D, with color schemes indicated for the modeled D273N, D273S, D273L in the WT IDH1 holo (1T0L (76)) and WT IDH1 apo (1T09 (76)) backgrounds. Previously solved structures of apo WT IDH1(76), holo IDH1(76), and C132H IDH1 (115) with ML309 docked (5K11 (115)) are also shown. We previously modeled the histidine mutation at residue 132 using a structure of R132C IDH1, and so denote this change as C132H IDH1. (B) Energy-minimized models of D273N, D273S, and D273L IDH1 generated in the WT IDH1 holo structure (αKG, NADP_+_, and Ca_2+_ bound, 1T0L (76)) aligned with apo (76) and holo (76) WT IDH1. (C) Energy-minimized models of D273N, D273S, and D273L IDH1 generated in the WT IDH1 apo structure (NADP_+_ bound, 1T09 (76)) aligned with apo (76) and holo (76) WT IDH1. (D) Energy-minimized models of D273N, D273S, D273L IDH1 generated in the WT IDH1 apo structure (NADP_+_ bound, 1T09 (76)) aligned with apo WT IDH1 apo (76) and C132H IDH1 with ML309 docked in the inhibitor binding pocket (115).

To extend our biochemical studies, we were interested in studying how changes in pH_i_ affected cellular αKG and isocitrate levels. Based on our kinetic data, our observation that HT1080 cells show a decrease in αKG levels upon a decrease in pH_i_ is unsurprising, though of course there are many sources of αKG production and consumption in addition to what is generated by IDH1. There are also a host of regulatory strategies to modulate levels of this metabolite. For example, in studies exploring the consequences of pH_i_ changes in porcine kidney cells, acidosis results in a decrease of αKG concentration, while alkylosis causes an increase (105). α-Ketoglutarate dehydrogenase is regulated by pH, with *K*_m_ values decreasing and *k*_cat_ values remaining relatively constant as the pH decreases. Thus, it is possible that acidic pH has a more potent effect on α-ketoglutarate dehydrogenase activity relative to IDH1 (106). To better isolate the cellular changes in isocitrate and αKG levels resulting from altered IDH1 activity upon a decrease in pH_i_, metabolic flux analysis studies in a series of IDH1 knockout and IDH1-amplified cell lines would be required (18,107).

While we focus in this work on the reactions catalyzed by WT IDH1, we do note that levels of 2HG decreased upon ESOM treatment. WT lactate dehydrogenase 1 and WT malate dehydrogenase 2 moonlighting activity of *L*-2HG production increases with decreasing pH (50,61), though production of D2HG by WT IDH1 does not appear to be dependent on pH (50). Thus, we were surprised to see a decrease in 2HG production. Of course, cell lines, especially cancer cell lines, have their own unique metabolic rewiring at play, and likely vary greatly in the regulation or production of 2HG pools. Furthermore, these metabolic addictions can change based on available nutrients, genetic profile, gene regulation, etc. An important future direction will be to assess the role of pH in regulating mutant IDH1 catalysis and resulting metabolite flux inside cells.

We also consider the possibility that treatment with proton pump inhibitors like ESOM can affect cells beyond simply lowering pH_i._ A possible consequence of ESOM treatment includes an increase in autophagy in some cell lines (108). We do note several amino acid levels are increased upon ESOM treatment (Table S1). Interestingly, some of the largest changes in amino acid composition are in the branched chain amino acids (L, I, V) in WT IDH1-expressing cell lines (Table S1). αKG and other α-keto acids are important for generating branched chain amino acids, and thus our observed depletion of αKG upon ESOM treatment may be related to increased branched chain amino acid synthesis (109).

Overall, our findings characterize the pH sensitivity of human IDH, showing that the *k*_cat_ of the forward reaction appears most sensitive to changes in pH. Moreover, we identify one residue, D273, that appears to sense these changes in pH by affecting catalysis. Critically, mutation of this residue ablates pH sensitivity. This work is important to further elucidate the mechanisms of human IDH1 catalysis, and clarify its regulation in distinct cellular microenvironments related to normal processes and disease. This work applies structural informatics and biochemical characterization to identify new residue clusters that play a novel functional role in regulating human IDH1 catalysis.

## EXPERIMENTAL PROCEDURES

### Materials

Tris-hydrochloric acid, tris base, bis-tris, NaOH, NaCl, MgCl_2_ hexahydrate, dithiothreitol, NADP_+_ disodium salt, NADPH, KPhos dibasic, KCl, BL-21 Gold (DE3) competent cells, Luria-broth (LB)-Agar, kanamycin sulfate, Terrific broth, IPTG, EDTA-free protease inhibitor tablets, Ni-NTA resin, Dulbecco’s Modified Eagle Medium (DMEM), 2’,7’-bis-(2-carboxyethyl)-5- (and-6)-carboxyfluorescein acetoxymethyl ester, Dulbecco’s phosphate buffered saline (DPBS) and Triton X-100 were all obtained from Fisher Scientific (Hampton, NH). Isocitrate, αKG, and imidazole were obtained from Acros Organics (Fisher Scientific, Hampton, NH). Fetal bovine serum (FBS) was obtained from VWR (Radnor, PA). 5-(*N*-Ethyl-*N*-isopropyl)amiloride was obtained from Sigma-Aldrich (St. Louis, MO). ESOM sodium salt was obtained from Apexbio (Houston, TX). Nigericin sodium was obtained from Tocris (Bristol, UK).

### Plasmid mutagenesis

WT and R132H IDH1 clones were generously provided by Charles Rock (St. Jude’s Hospital). All IDH1 cDNA constructs are in a pET-28b plasmid vector that contains an *N*-terminal hexahistidine tag. Site-directed mutagenesis was used to generate single point mutations in WT IDH1 using the manufacturer’s (Kappa Biosciences, Oslo, Norway) guidelines with the following primers: D273N (forward primer 5’-TTTATTTGGGCCTGCAAAAACTATAATGGTGATGTTCAGAGC, reverse primer 5’-GCTCTGAACATCACCATTATAGTTTTTGCAGGCCCAAATAAA); D273L (forward primer 5’-TGGTTTTATTTGGGCCTGCAAAAACTATCTGGGTGATGTTCAGAGCGA, reverse primer 5’-TCGCTCTGAACATCACCCAGATAGTTTTTGCAGGCCCAAATAAAACCA); D273S (forward primer 5’-GTGGTTTTATTTGGGCCTGCAAAAACTATAGTGGTGATGTTCAGAG, reverse primer 5’-CTCTGAACATCACCACTATAGTTTTTGCAGGCCCAAATAAAACCAC); K217M (forward primer 5’-CTGAGCACCAAAAATACCATTCTGATGAAATACGATGGTCGCTTTAAAGATA, reverse primer 5’-TATCTTTAAAGCGACCATCGTATTTCATCAGAATGGTATTTTTGGTGCTCAG); K217Q (forward 5’-CTGAGCACCAAAAATACCATTCTGCAGAAATACGATGGTCGCTTTAAAGAT, reverse primer 5’-ATCTTTAAAGCGACCATCGTATTTCTGCAGAATGGTATTTTTGGTGCTCAG). Site-directed mutagenesis was also used to generate single point mutations in the R132H IDH1 background according to the manufacturer’s guidelines (Kappa Biosciences, Oslo, Norway) using the following primers: D273N (forward primer 5’-TTTATTTGGGCCTGCAAAAACTATAATGGTGATGTTCAGAGC, reverse primer 5’-GCTCTGAACATCACCATTATAGTTTTTGCAGGCCCAAATAAA); D273L (forward primer 5’-TGGTTTTATTTGGGCCTGCAAAAACTATCTGGGTGATGTTCAGAGCGA, reverse primer 5’-TCGCTCTGAACATCACCCAGATAGTTTTTGCAGGCCCAAATAAAACCA); D273S (forward primer 5’-GTGGTTTTATTTGGGCCTGCAAAAACTATAGTGGTGATGTTCAGAG, reverse primer 5’-CTCTGAACATCACCACTATAGTTTTTGCAGGCCCAAATAAAACCAC). All constructs were sequenced to confirm accuracy.

### Protein purification

WT and mutant IDH1 homodimers were expressed and purified as described previously (> 95% purity) (110). Briefly, BL21-Gold (DE3) *E. coli* cells transformed with the proper IDH1 construct were incubated at 37 °C to an OD_600_ of 1.0-1.2. The cultures were then induced with a final concentration of 1 mM IPTG and incubated for 18-20 hours at 18 °C. IDH1 protein was purified using Ni-NTA column chromatography, and flash frozen in liquid nitrogen for storage at -80°C for ≤1 month.

### Steady-state activity assays

The activity of IDH1 homodimers was assessed at 37 °C using steady-state methods described previously (110) (62). The tris-based assays were prepared as follows: for the conversion of isocitrate to αKG for pH >7, a cuvette containing assay buffer (50 mM tris buffer ranging from pH 8.0 to 7.2 at 37°C, 150 mM NaCl, 10 mM MgCl_2_, 0.1 mM dithiothreitol), and 100 nM WT IDH1, were preincubated for 3 minutes at 37 °C. For the conversion of isocitrate to αKG for pH ≤7.0 a cuvette containing assay buffer (50 mM bis-tris ranging from pH 7.0 to 6.2 at 37°C, 150 mM 150 NaCl, 10 mM MgCl_2_, 0.1 mM dithiothreitol), and 100 nM WT IDH1 were preincubated for 3 minutes at 37 °C. For the conversion of αKG to isocitrate for pH >7 (62), a cuvette containing assay buffer (50 mM tris ranging from pH 8.0-7.2 at 37 °C, 150 mM NaCl, 10 mM MgCl_2_, 66 mM NaHCO_3_), and 100 nM WT IDH1, were preincubated for 3 minutes at 37 °C. For the conversion of αKG to isocitrate for pH ≤7.0, a cuvette containing assay buffer (50 mM bis-tris pH ranging 7.0-6.2 for 37°C, 150 mM NaCl, 10 mM MgCl_2_, 66 mM NaHCO_3_) and 100 nM WT IDH1, were preincubated for 3 minutes at 37 °C.

The KPhos buffer-based assays were prepared as follows: for the conversion of isocitrate to αKG, a cuvette containing assay buffer (50 mM KPhos ranging from pH 8.0-6.2 at 37 °C, 150 mM KCl, 10 MgCl_2_, 0.1 mM dithiotheitol), and 100 nM IDH1 were preincubated for 3 minutes at 37°C. For the conversion of αKG to isocitrate, a cuvette containing assay buffer (50 mM KPhos pH ranging from 8.0-6.2 at 37 °C, 150 mM KCl, 10 mM MgCl_2_, 66 mM NaHCO_3_, and 0.1 mM dithiothreitol), and 100 nM IDH1, were preincubated for 3 minutes at 37 °C.

The pH of αKG was adjusted to the pH of the assay before use. In reactions measuring isocitrate to αKG conversion, reactions were initiated by adding NADP_+_ and isocitrate, with saturating NADP_+_ and titrating in various concentrations of isocitrate. In reactions measuring αKG to isocitrate conversion, or αKG to D2HG conversion, reactions were initiated by adding NADPH and αKG, with saturating NADPH and titrating in various concentrations of αKG. In all cases, the change in absorbance due to changing NADPH concentrations was monitored at 340 nm.

For all reactions, the slope of the linear ranges of the assays were calculated and converted to µM NADPH using the molar extinction coefficient for NADPH of 6.22 cm_-1_ mM_-1_ to obtain *k*_obs_ (i.e. µM NADPH/µM enzyme s_-1_ at each substrate concentration). Results were fit to hyperbolic plots in GraphPad Prism (GraphPad Software, La Jolla, CA) to estimate *k*_*cat*_ and *K*_m_ values ± SE.

### Thermal stability using circular dichroism

The stability of WT IDH1 in the pH range from 8.0-6.2 was assessed using circular dichroism as described previously (110). Briefly, WT IDH1 was diluted to 5 μM in a buffer containing 10 mM KPhos buffer at desired pH and 100 mM KCl. The thermal melt experiment was initiated at 20 °C and the temperature was increased to 70 °C in 1°C increments. The secondary structure of IDH1 is rich in alpha helices and was monitored via the 222 nm peak, corresponding to α-helicity. Analysis was performed using GraphPad Prism (GraphPad Software, La Jolla, CA) with a Boltzmann sigmoidal fit (111).

### pHinder algorithm

The details of the pHinder algorithm have been described previously (29,43,74). Briefly, protein structures were downloaded from the Protein Data Bank (PDB) and the pHinder program was used to calculate ionizable residue networks using a two-step procedure. First, a Delaunay triangulation was calculated using the terminal side-chain atoms of all ionizable residues (D, E, H, C, K, R) and minimized by removing network edges longer than 10 Å. This triangulation was further simplified by removing redundant network connections. Second, using a molecular surface calculated by the pHinder algorithm, each ionizable network residue was classified as buried (>3.0 Å below the surface), margin (<3.0 Å below and <1.0 Å above the surface), or exposed (>1.0 Å above the surface). Depth of burial was determined by measuring the minimum distance between the ionizable group and the triangular facets of the pHinder-calculated surface. Buried networks were identified as contiguous runs of buried residues. Acidic and basic networks were identified as contiguous runs of each residue type.

### ITC measurements

ITC was performed in Sanford Burnham Prebys Protein Analysis Core using a Low Volume Affinity ITC calorimeter (TA Instruments). 3.0 to 6.0 μl aliquots of solution containing 0.15 mM AGI-5198 or ML309 were injected into the cell containing 0.03 to 0.05 mM protein. 20 injections were made. The experiments were performed at 23 °C in buffer containing 20 mM Tris pH 7.5, 100 mM NaCl, 2 mM β-mercaptoethanol, and 0.2 mM NADPH. Baseline control data were collected injecting ligand into the cell containing buffer only. ITC data were analyzed using Nanoanalyze software provided by TA Instruments.

### Computational methods

PROPKA (77,78) was used to predict the protonation states in the PDB 1T0L structure (76) as described previously. For basic structural modeling of the mutants, due to structural similarity in the mutant and WT IDH1 holo forms, and instability of the α10 helix in the mutant IDH1 structures, the WT background was selected for basic modeling. PDB 1T0L and 1T09 structures (76) were used to generate models of D273N, D273L, and D273S in the WT IDH1 apo (NADP_+_ bound) and holo (isocitrate, NADP_+_, and Ca_2+_) backgrounds. Mutations were made using Coot software (112), and Geometry Minimization in the Phenix software suite (113) was used to regularize geometries of the ligands and protein chains. 500 maximum iterations and 5 macrocycles were used, with bond lengths, bond angles, nonbonded distances, dihedral angles, chirality, parallelity, and planarity considered.

### Cellular pH_i_ modulations

HT1080 cell lines were all cultured in DMEM with 10% FBS and incubated at 37 °C with 5% CO_2_. The effects of proton pump inhibitors on pH_i_ was evaluated in triplicate by first adding 0.3×10_5_ cells per well in a 24-well plate and incubated overnight. The following day the proton pump inhibitor ESOM was dissolved in PBS and was added to the cells (200 µM final concentration), and the cells were again incubated at 37°C 5% CO_2_. After 16-24 hours, the pH_i_ was read by first loading cells with 1 μM BCECF-AM dissolved in DMSO (final cellular concentration of DMSO at 0.1%) at 37 °C at 5% CO_2_ for 15 minutes. Excess dye was removed by washing cells 3 times with bicarbonate buffer (25 mM bicarbonate, 115 mM NaCl, 5 mM KCl, 10 mM glucose, 1 mM MgSO_4_, 1 mM KHPO_4_ pH 7.4 at 37 °C, and 2 mM CaCl_2_) with ESOM to preserve pH_i_ manipulation conditions. After wash steps, the fluorescence intensity of the BCECF-AM probe was detected with dual excitation (490 nm and 440 nm) and a fixed emission wavelength of 535 nm. Fluorescence ratios (490 nm/440 nm) were converted to pH_i_ values by calibrating each experiment with a standard curve ranging from pH 6.5 to 7.5 using 25 mM HEPES, 105 mM KCl, and 1 mM MgCl_2_ and 10 μM Nigericin to equilibrate pH_i_ with pH_e_. Metabolites were derivatized and quantified using GC-MS as described previously (114).

## Acknowledgments

The authors would like to thank Prof. Kun-Liang Guan for the HT1080 cell lines. This work was funded by a Research Scholar Grant, RSG-19-075-01-TBE, from the American Cancer Society (CDS), National Institutes of Health R00 CA187594 (CDS), R01 CA197855 (DLB), SBP NCI Cancer Center Support Grant P30 CA030199 (OZ and DAS), National Institutes of Health R35GM119518, R01NS103906, and R03TR002908 (DGI), U54CA132384 (SDSU) & U54CA132379 (UC San Diego), MARC 5T34GM008303 (SDSU), and IMSD 5R25GM058906 (SDSU), and the California Metabolic Research Foundation (SDSU). The results shown here are in part based upon data generated by the TCGA Research Network: https://www.cancer.gov/tcga. The content is solely the responsibility of the authors and does not necessarily represent the official views of the National Institutes of Health.

## Author contributions

LAL and ZL performed the kinetic assays. LAL prepared cDNA constructs and performed the cellular pH modulation assays and CD measurements. KAW and DLB aided in the design of the cellular pH modulation assays and provided experimental training. DAS and OZ performed the cellular metabolite quantitation. JMS ran the calculations predicting residue pK_a_ values and performed the ML309 docking. AH aided in protein preparations and cellular studies. AAB performed the ITC experiments. DGI performed the pHinder algorithm and analyzed the results. CDS conceived the idea for the project, analyzed the pHinder algorithm results, performed the energy minimizations, and wrote the paper. All contributed to data analysis and manuscript editing and approved the final version of the manuscript.

## FOOTNOTES

## The abbreviations used are

CD: circular dichroism;
*E. coli*: *Escherichia coli*;
DIDS: 4,4’-diisothiocyano-2,2’-stilbenedisulfonic acid;
DMEM: Dulbecco’s Modified Eagle Medium;
DPBS: Dulbecco’s phosphate buffered saline;
EIPA: 5-(*N*-ethyl-*N*-isopropyl)amiloride;
ESOM: esomeprazole;
FBS: Fetal bovine serum;
GTP: guanosine triphosphate;
IDH: isocitrate dehydrogenase;
IPTG: isopropyl β-d-1-thiogalactopyranoside;
αKG: α-ketoglutarate;
KPhos: potassium phosphate;
NADPH: ß-nicotinamide adenine dinucleotide phosphate reduced;
NADP_+_: ß-nicotinamide adenine dinucleotide phosphate;
N.D.: not detected;
ns: non-significant;
PDB: protein databank;
pH_i_: internal pH;
pH_e_: external pH;
S.E.M: standard error of the mean;
WT: wild type;
GC/MS: gas chromatography/mass spectrometry

## REFERENCES

1. Mailloux, R. J., Beriault, R., Lemire, J., Singh, R., Chenier, D. R., Hamel, R. D., and Appanna, V. D. (2007) The tricarboxylic acid cycle, an ancient metabolic network with a novel twist. PLoS One 2, e690

2. Reitman, Z. J., and Yan, H. (2010) Isocitrate dehydrogenase 1 and 2 mutations in cancer: alterations at a crossroads of cellular metabolism. J Natl Cancer Inst 102, 932–941

3. Bogdanovic, E. (2015) IDH1, lipid metabolism and cancer: Shedding new light on old ideas. Biochim Biophys Acta 1850, 1781–1785

4. Calvert, A. E., Chalastanis, A., Wu, Y., Hurley, L. A., Kouri, F. M., Bi, Y., Kachman, M., May, J. L., Bartom, E., Hua, Y., Mishra, R. K., Schiltz, G. E., Dubrovskyi, O., Mazar, A. P., Peter, M. E., Zheng, H., James, C. D., Burant, C. F., Chandel, N. S., Davuluri, R. V., Horbinski, C., and Stegh, A. H. (2017) Cancer-associated IDH1 promotes growth and resistance to targeted therapies in the absence of mutation. Cell Rep 19, 1858–1873

5. Cerami, E., Gao, J., Dogrusoz, U., Gross, B. E., Sumer, S. O., Aksoy, B. A., Jacobsen, A., Byrne, C. J., Heuer, M. L., Larsson, E., Antipin, Y., Reva, B., Goldberg, A. P., Sander, C., and Schultz, N. (2012) The cBio cancer genomics portal: an open platform for exploring multidimensional cancer genomics data. Cancer Discov 2, 401–404

6. Tokeshi, Y., Shimada, S., Hanashiro, K., Sunagawa, M., Nakamura, M., and Kosugi, T. (2001) The nucleotide sequence of dinitrophenyl-specific IgE and Fc(epsilon)RI alpha-subunit obtained from FE-3 hybridoma cells. Hybrid Hybridomics 20, 361–368

7. Parsons, D. W., Jones, S., Zhang, X., Lin, J. C., Leary, R. J., Angenendt, P., Mankoo, P., Carter, H., Siu, I. M., Gallia, G. L., Olivi, A., McLendon, R., Rasheed, B. A., Keir, S., Nikolskaya, T., Nikolsky, Y., Busam, D. A., Tekleab, H., Diaz, L. A., Jr., Hartigan, J., Smith, D. R., Strausberg, R. L., Marie, S. K., Shinjo, S. M., Yan, H., Riggins, G. J., Bigner, D. D., Karchin, R., Papadopoulos, N., Parmigiani, G., Vogelstein, B., Velculescu, V. E., and Kinzler, K. W. (2008) An integrated genomic analysis of human glioblastoma multiforme. Science 321, 1807–1812

8. Yan, H., Parsons, D. W., Jin, G., McLendon, R., Rasheed, B. A., Yuan, W., Kos, I., Batinic-Haberle, I., Jones, S., Riggins, G. J., Friedman, H., Friedman, A., Reardon, D., Herndon, J., Kinzler, K. W., Velculescu, V. E., Vogelstein, B., and Bigner, D. D. (2009) IDH1 and IDH2 mutations in gliomas. N Engl J Med 360, 765–773

9. Mardis, E. R., Ding, L., Dooling, D. J., Larson, D. E., McLellan, M. D., Chen, K., Koboldt, D. C., Fulton, R. S., Delehaunty, K. D., McGrath, S. D., Fulton, L. A., Locke, D. P., Magrini, V. J., Abbott, R. M., Vickery, T. L., Reed, J. S., Robinson, J. S., Wylie, T., Smith, S. M., Carmichael, L., Eldred, J. M., Harris, C. C., Walker, J., Peck, J. B., Du, F., Dukes, A. F., Sanderson, G. E., Brummett, A. M., Clark, E., McMichael, J. F., Meyer, R. J., Schindler, J. K., Pohl, C. S., Wallis, J. W., Shi, X., Lin, L., Schmidt, H., Tang, Y., Haipek, C., Wiechert, M. E., Ivy, J. V., Kalicki, J., Elliott, G., Ries, R. E., Payton, J. E., Westervelt, P., Tomasson, M. H., Watson, M. A., Baty, J., Heath, S., Shannon, W. D., Nagarajan, R., Link, D. C., Walter, M. J., Graubert, T. A., DiPersio, J. F., Wilson, R. K., and Ley, T. J. (2009) Recurring mutations found by sequencing an acute myeloid leukemia genome. N Engl J Med 361, 1058–1066

10. Kosmider, O., Gelsi-Boyer, V., Slama, L., Dreyfus, F., Beyne-Rauzy, O., Quesnel, B., Hunault-Berger, M., Slama, B., Vey, N., Lacombe, C., Solary, E., Birnbaum, D., Bernard, O. A., and Fontenay, M. (2010) Mutations of IDH1 and IDH2 genes in early and accelerated phases of myelodysplastic syndromes and MDS/myeloproliferative neoplasms. Leukemia 24, 1094–1096

11. Borger, D. R., Tanabe, K. K., Fan, K. C., Lopez, H. U., Fantin, V. R., Straley, K. S., Schenkein, D. P., Hezel, A. F., Ancukiewicz, M., Liebman, H. M., Kwak, E. L., Clark, J. W., Ryan, D. P., Deshpande, V., Dias-Santagata, D., Ellisen, L. W., Zhu, A. X., and Iafrate, A. J. (2012) Frequent mutation of isocitrate dehydrogenase (IDH)1 and IDH2 in cholangiocarcinoma identified through broad-based tumor genotyping. Oncologist 17, 72–79

12. Pietrak, B., Zhao, H., Qi, H., Quinn, C., Gao, E., Boyer, J. G., Concha, N., Brown, K., Duraiswami, C., Wooster, R., Sweitzer, S., and Schwartz, B. (2011) A tale of two subunits: how the neomorphic R132H IDH1 mutation enhances production of alphaHG. Biochemistry 50, 4804–4812

13. Dang, L., White, D. W., Gross, S., Bennett, B. D., Bittinger, M. A., Driggers, E. M., Fantin, V. R., Jang, H. G., Jin, S., Keenan, M. C., Marks, K. M., Prins, R. M., Ward, P. S., Yen, K. E., Liau, L. M., Rabinowitz, J. D., Cantley, L. C., Thompson, C. B., Vander Heiden, M. G., and Su, S. M. (2009) Cancer-associated IDH1 mutations produce 2-hydroxyglutarate. Nature 462, 739–744

14. Dang, L., Jin, S., and Su, S. M. (2010) IDH mutations in glioma and acute myeloid leukemia. Trends Mol Med 16, 387–397

15. Horbinski, C. (2013) What do we know about IDH1/2 mutations so far, and how do we use it? Acta Neuropathol 125, 621–636

16. Popovici-Muller, J., Saunders, J. O., Salituro, F. G., Travins, J. M., Yan, S., Zhao, F., Gross, S., Dang, L., Yen, K. E., Yang, H., Straley, K. S., Jin, S., Kunii, K., Fantin, V. R., Zhang, S., Pan, Q., Shi, D., Biller, S. A., and Su, S. M. (2012) Discovery of the first potent inhibitors of mutant IDH1 that lower tumor 2-HG in vivo. ACS Med Chem Lett 3, 850–855

17. DiNardo, C. D., Stein, E. M., de Botton, S., Roboz, G. J., Altman, J. K., Mims, A. S., Swords, R., Collins, R. H., Mannis, G. N., Pollyea, D. A., Donnellan, W., Fathi, A. T., Pigneux, A., Erba, H. P., Prince, G. T., Stein, A. S., Uy, G. L., Foran, J. M., Traer, E., Stuart, R. K., Arellano, M. L., Slack, J. L., Sekeres, M. A., Willekens, C., Choe, S., Wang, H., Zhang, V., Yen, K. E., Kapsalis, S. M., Yang, H., Dai, D., Fan, B., Goldwasser, M., Liu, H., Agresta, S., Wu, B., Attar, E. C., Tallman, M. S., Stone, R. M., and Kantarjian, H. M. (2018) Durable remissions with Ivosidenib in IDH1-mutated relapsed or refractory AML. N Engl J Med 378, 2386–2398

18. Metallo, C. M., Gameiro, P. A., Bell, E. L., Mattaini, K. R., Yang, J., Hiller, K., Jewell, C. M., Johnson, Z. R., Irvine, D. J., Guarente, L., Kelleher, J. K., Vander Heiden, M. G., Iliopoulos, O., and Stephanopoulos, G. (2012) Reductive glutamine metabolism by IDH1 mediates lipogenesis under hypoxia. Nature 481, 380–384

19. Wise, D. R., Ward, P. S., Shay, J. E., Cross, J. R., Gruber, J. J., Sachdeva, U. M., Platt, J. M., DeMatteo, R. G., Simon, M. C., and Thompson, C. B. (2011) Hypoxia promotes isocitrate dehydrogenase-dependent carboxylation of alpha-ketoglutarate to citrate to support cell growth and viability. Proc Natl Acad Sci U.S.A 108, 19611–19616

20. Filipp, F. V., Scott, D. A., Ronai, Z. A., Osterman, A. L., and Smith, J. W. (2012) Reverse TCA cycle flux through isocitrate dehydrogenases 1 and 2 is required for lipogenesis in hypoxic melanoma cells. Pigment Cell Melanoma Res 25, 375–383

21. Seltzer, M. J., Bennett, B. D., Joshi, A. D., Gao, P., Thomas, A. G., Ferraris, D. V., Tsukamoto, T., Rojas, C. J., Slusher, B. S., Rabinowitz, J. D., Dang, C. V., and Riggins, G. J. (2010) Inhibition of glutaminase preferentially slows growth of glioma cells with mutant IDH1. Cancer Res 70, 8981–8987

22. Grassian, A. R., Parker, S. J., Davidson, S. M., Divakaruni, A. S., Green, C. R., Zhang, X., Slocum, K. L., Pu, M., Lin, F., Vickers, C., Joud-Caldwell, C., Chung, F., Yin, H., Handly, E. D., Straub, C., Growney, J. D., Vander Heiden, M. G., Murphy, A. N., Pagliarini, R., and Metallo, C. M. (2014) IDH1 mutations alter citric acid cycle metabolism and increase dependence on oxidative mitochondrial metabolism. Cancer Res 74, 3317–3331

23. Reitman, Z. J., Duncan, C. G., Poteet, E., Winters, A., Yan, L. J., Gooden, D. M., Spasojevic, I., Boros, L. G., Yang, S. H., and Yan, H. (2014) Cancer-associated isocitrate dehydrogenase 1 (IDH1) R132H mutation and D-2-Hydroxyglutarate stimulate glutamine metabolism under hypoxia. J Biol Chem 289, 23318–23328

24. Mullen, A. R., Hu, Z., Shi, X., Jiang, L., Boroughs, L. K., Kovacs, Z., Boriack, R., Rakheja, D., Sullivan, L. B., Linehan, W. M., Chandel, N. S., and DeBerardinis, R. J. (2014) Oxidation of alpha-ketoglutarate is required for reductive carboxylation in cancer cells with mitochondrial defects. Cell Rep 7, 1679–1690

25. Schonichen, A., Webb, B. A., Jacobson, M. P., and Barber, D. L. (2013) Considering protonation as a posttranslational modification regulating protein structure and function. Annu Rev Biophys 42, 289–314

26. Matsuyama, S., Llopis, J., Deveraux, Q. L., Tsien, R. Y., and Reed, J. C. (2000) Changes in intramitochondrial and cytosolic pH: early events that modulate caspase activation during apoptosis. Nat Cell Biol 2, 318–325

27. Ciriolo, M. R., Palamara, A. T., Incerpi, S., Lafavia, E., Bue, M. C., De Vito, P., Garaci, E., and Rotilio, G. (1997) Loss of GSH, oxidative stress, and decrease of intracellular pH as sequential steps in viral infection. J Biol Chem 272, 2700–2708

28. Isom, D. G., Page, S. C., Collins, L. B., Kapolka, N. J., Taghon, G. J., and Dohlman, H. G. (2018) Coordinated regulation of intracellular pH by two glucose-sensing pathways in yeast. J Biol Chem 293, 2318–2329

29. Isom, D. G., Sridharan, V., Baker, R., Clement, S. T., Smalley, D. M., and Dohlman, H. G. (2013) Protons as second messenger regulators of G protein signaling. Mol Cell 51, 531–538

30. Mulkey, D. K., Henderson, R. A., 3rd, Ritucci, N. A., Putnam, R. W., and Dean, J. B. (2004) Oxidative stress decreases pHi and Na(+)/H(+) exchange and increases excitability of solitary complex neurons from rat brain slices. Am J Physiol Cell Physiol 286, C940–951

31. Nakamura, U., Iwase, M., Uchizono, Y., Sonoki, K., Sasaki, N., Imoto, H., Goto, D., and Iida, M. (2006) Rapid intracellular acidification and cell death by H2O2 and alloxan in pancreatic beta cells. Free Radic Biol Med 40, 2047–2055

32. Ulmschneider, B., Grillo-Hill, B. K., Benitez, M., Azimova, D. R., Barber, D. L., and Nystul, T. G. (2016) Increased intracellular pH is necessary for adult epithelial and embryonic stem cell differentiation. J Cell Biol 215, 345–355

33. Denker, S. P., and Barber, D. L. (2002) Cell migration requires both ion translocation and cytoskeletal anchoring by the Na-H exchanger NHE1. J Cell Biol 159, 1087–1096

34. Harguindey, S., Reshkin, S. J., Orive, G., Arranz, J. L., and Anitua, E. (2007) Growth and trophic factors, pH and the Na+/H+ exchanger in Alzheimer’s disease, other neurodegenerative diseases and cancer: new therapeutic possibilities and potential dangers. Curr Alzheimer Res 4, 53–65

35. Tracz, S. M., Abedini, A., Driscoll, M., and Raleigh, D. P. (2004) Role of aromatic interactions in amyloid formation by peptides derived from human Amylin. Biochemistry 43, 15901–15908

36. White, K. A., Ruiz, D. G., Szpiech, Z. A., Strauli, N. B., Hernandez, R. D., Jacobson, M. P., and Barber, D. L. (2017) Cancer-associated arginine-to-histidine mutations confer a gain in pH sensing to mutant proteins. Sci Signal 10

37. Webb, B. A., Chimenti, M., Jacobson, M. P., and Barber, D. L. (2011) Dysregulated pH: a perfect storm for cancer progression. Nat Rev Cancer 11, 671–677

38. Corbet, C., and Feron, O. (2017) Tumour acidosis: from the passenger to the driver’s seat. Nat Rev Cancer 17, 577–593

39. Swietach, P., Vaughan-Jones, R. D., Harris, A. L., and Hulikova, A. (2014) The chemistry, physiology and pathology of pH in cancer. Philos Trans R Soc Lond B Biol Sci 369, 20130099

40. Karp, D. A., Stahley, M. R., and Garcia-Moreno, B. (2010) Conformational consequences of ionization of Lys, Asp, and Glu buried at position 66 in staphylococcal nuclease. Biochemistry 49, 4138–4146

41. Bell-Upp, P., Robinson, A. C., Whitten, S. T., Wheeler, E. L., Lin, J., Stites, W. E., and E, B. G. (2011) Thermodynamic principles for the engineering of pH-driven conformational switches and acid insensitive proteins. Biophys Chem 159, 217–226

42. Isom, D. G., Castaneda, C. A., Cannon, B. R., and Garcia-Moreno, B. (2011) Large shifts in pKa values of lysine residues buried inside a protein. Proc Natl Acad Sci U.S.A 108, 5260–5265

43. Isom, D. G., Sridharan, V., and Dohlman, H. G. (2016) Regulation of Ras paralog thermostability by networks of buried ionizable groups. Biochemistry 55, 534–542

44. Ludwig, M. G., Vanek, M., Guerini, D., Gasser, J. A., Jones, C. E., Junker, U., Hofstetter, H., Wolf, R. M., and Seuwen, K. (2003) Proton-sensing G-protein-coupled receptors. Nature 425, 93–98

45. Webb, B. A., White, K. A., Grillo-Hill, B. K., Schonichen, A., Choi, C., and Barber, D. L. (2016) A histidine cluster in the cytoplasmic domain of the Na-H exchanger NHE1 confers pH-sensitive phospholipid binding and regulates transporter activity. J Biol Chem 291, 24096–24104

46. White, K. A., Grillo-Hill, B. K., Esquivel, M., Peralta, J., Bui, V. N., Chire, I., and Barber, D. L. (2018) beta-Catenin is a pH sensor with decreased stability at higher intracellular pH. J Cell Biol 217, 3965–3976

47. Choi, C. H., Webb, B. A., Chimenti, M. S., Jacobson, M. P., and Barber, D. L. (2013) pH sensing by FAK-His58 regulates focal adhesion remodeling. J Cell Biol 202, 849–859

48. Vercoulen, Y., Kondo, Y., Iwig, J. S., Janssen, A. B., White, K. A., Amini, M., Barber, D. L., Kuriyan, J., and Roose, J. P. (2017) A histidine pH sensor regulates activation of the Ras-specific guanine nucleotide exchange factor RasGRP1. eLife 6

49. Corbet, C., Pinto, A., Martherus, R., Santiago de Jesus, J. P., Polet, F., and Feron, O. (2016) Acidosis Drives the Reprogramming of Fatty Acid Metabolism in Cancer Cells through Changes in Mitochondrial and Histone Acetylation. Cell Metab 24, 311–323

50. Intlekofer, A. M., Wang, B., Liu, H., Shah, H., Carmona-Fontaine, C., Rustenburg, A. S., Salah, S., Gunner, M. R., Chodera, J. D., Cross, J. R., and Thompson, C. B. (2017) L-2-Hydroxyglutarate production arises from noncanonical enzyme function at acidic pH. Nat Chem Biol 13, 494–500

51. Bailey, J., Bell, E. T., and Bell, J. E. (1982) Regulation of bovine glutamate dehydrogenase. The effects of pH and ADP. J Biol Chem 257, 5579–5583

52. Nissim, I. (1999) Newer aspects of glutamine/glutamate metabolism: the role of acute pH changes. Am J Physiol 277, F493–497

53. Tannen, R. L., and Kunin, A. S. (1981) Effect of pH on metabolism of alpha-ketoglutarate by renal cortical mitochondria. Am J Physiol 240, F120–126

54. Wood, D. C., Hodges, C. T., Howell, S. M., Clary, L. G., and Harrison, J. H. (1981) The N-ethylmaleimide-sensitive cysteine residue in the pH-dependent subunit interactions of malate dehydrogenase. J Biol Chem 256, 9895–9900

55. Trivedi, B., and Danforth, W. H. (1966) Effect of pH on the kinetics of frog muscle phosphofructokinase. J Biol Chem 241, 4110–4112

56. Casey, J. R., Grinstein, S., and Orlowski, J. (2010) Sensors and regulators of intracellular pH. Nat Rev Mol Cell Biol 11, 50–61

57. Webb, B. A., Forouhar, F., Szu, F. E., Seetharaman, J., Tong, L., and Barber, D. L. (2015) Structures of human phosphofructokinase-1 and atomic basis of cancer-associated mutations. Nature 523, 111–114

58. Gaspar, P., Neves, A. R., Shearman, C. A., Gasson, M. J., Baptista, A. M., Turner, D. L., Soares, C. M., and Santos, H. (2007) The lactate dehydrogenases encoded by the ldh and ldhB genes in *Lactococcus lactis* exhibit distinct regulation and catalytic properties-comparative modeling to probe the molecular basis. FEBS J 274, 5924–5936

59. Suzuki, H., and Ogura, Y. (1970) Effect of pH on the kinetic parameters of yeast L(+)-lactate dehydrogenase (cytochrome b2). J Biochem 67, 291–295

60. Dobson, G. P., Yamamoto, E., and Hochachka, P. W. (1986) Phosphofructokinase control in muscle: nature and reversal of pH-dependent ATP inhibition. Am J Physiol 250, R71–76

61. Nadtochiy, S. M., Schafer, X., Fu, D., Nehrke, K., Munger, J., and Brookes, P. S. (2016) Acidic pH is a metabolic switch for 2-hydroxyglutarate generation and signaling. J Biol Chem 291, 20188–20197

62. Leonardi, R., Subramanian, C., Jackowski, S., and Rock, C. O. (2012) Cancer-associated isocitrate dehydrogenase mutations inactivate NADPH-dependent reductive carboxylation. J Biol Chem 287, 14615–14620

63. Huang, Y. C., and Colman, R. F. (2002) Evaluation by mutagenesis of the roles of His309, His315, and His319 in the coenzyme site of pig heart NADP-dependent isocitrate dehydrogenase. Biochemistry 41, 5637–5643

64. Rendina, A. R., Pietrak, B., Smallwood, A., Zhao, H., Qi, H., Quinn, C., Adams, N. D., Concha, N., Duraiswami, C., Thrall, S. H., Sweitzer, S., and Schwartz, B. (2013) Mutant IDH1 enhances the production of 2-hydroxyglutarate due to its kinetic mechanism. Biochemistry 52, 4563–4577

65. Vinekar, R., Verma, C., and Ghosh, I. (2012) Functional relevance of dynamic properties of Dimeric NADP-dependent Isocitrate Dehydrogenases. BMC Bioinformatics 13 Suppl 17, S2

66. Dalziel, K., and Londesborough, J. C. (1968) The mechanisms of reductive carboxylation reactions. Carbon dioxide or bicarbonate as substrate of nicotinamide-adenine dinucleotide phosphate-linked isocitrate dehydrogenase and malic enzyme. Biochem J 110, 223–230

67. Swenson, E. R. (2016) Hypoxia and Its Acid-Base Consequences: From Mountains to Malignancy. Adv Exp Med Biol 903, 301–323

68. Baldini, N., and Avnet, S. (2018) The Effects of Systemic and Local Acidosis on Insulin Resistance and Signaling. Int J Mol Sci 20

69. Grillo-Hill, B. K., Webb, B. A., and Barber, D. L. (2014) Ratiometric imaging of pH probes. Methods Cell Biol 123, 429–448

70. Srivastava, J., Barber, D. L., and Jacobson, M. P. (2007) Intracellular pH sensors: design principles and functional significance. Physiology (Bethesda) 22, 30–39

71. Ma, S., Jiang, B., Deng, W., Gu, Z. K., Wu, F. Z., Li, T., Xia, Y., Yang, H., Ye, D., Xiong, Y., and Guan, K. L. (2015) D-2-hydroxyglutarate is essential for maintaining oncogenic property of mutant IDH-containing cancer cells but dispensable for cell growth. Oncotarget 6, 8606–8620

72. Davis, M. I., Gross, S., Shen, M., Straley, K. S., Pragani, R., Lea, W. A., Popovici-Muller, J., Delabarre, B., Artin, E., Thorne, N., Auld, D. S., Li, Z., Dang, L., Boxer, M. B., and Simeonov, A. (2014) Biochemical, cellular and biophysical characterization of a potent inhibitor of mutant isocitrate dehydrogenase IDH1. J Biol Chem 289, 13717–13725

73. Jin, G., Pirozzi, C. J., Chen, L. H., Lopez, G. Y., Duncan, C. G., Feng, J., Spasojevic, I., Bigner, D. D., He, Y., and Yan, H. (2012) Mutant IDH1 is required for IDH1 mutated tumor cell growth. Oncotarget 3, 774–782

74. Isom, D. G., and Dohlman, H. G. (2015) Buried ionizable networks are an ancient hallmark of G protein-coupled receptor activation. Proc Natl Acad Sci U S A 112, 5702–5707

75. Harms, M. J., Castaneda, C. A., Schlessman, J. L., Sue, G. R., Isom, D. G., Cannon, B. R., and Garcia-Moreno, E. B. (2009) The pK(a) values of acidic and basic residues buried at the same internal location in a protein are governed by different factors. J Mol Biol 389, 34–47

76. Xu, X., Zhao, J., Xu, Z., Peng, B., Huang, Q., Arnold, E., and Ding, J. (2004) Structures of human cytosolic NADP-dependent isocitrate dehydrogenase reveal a novel self-regulatory mechanism of activity. J Biol Chem 279, 33946–33957

77. Sastry, G. M., Adzhigirey, M., Day, T., Annabhimoju, R., and Sherman, W. (2013) Protein and ligand preparation: parameters, protocols, and influence on virtual screening enrichments. J Comput Aided Mol Des 27, 221–234

78. Schrödinger Release 2018-3, Schrödinger Suite 2018-3 Protein Preparation Wizard.

79. Harms, M. J., Schlessman, J. L., Sue, G. R., and Garcia-Moreno, B. (2011) Arginine residues at internal positions in a protein are always charged. Proc Natl Acad Sci U.S.A 108, 18954–18959

80. Xie, X., Baird, D., Bowen, K., Capka, V., Chen, J., Chenail, G., Cho, Y., Dooley, J., Farsidjani, A., Fortin, P., Kohls, D., Kulathila, R., Lin, F., McKay, D., Rodrigues, L., Sage, D., Toure, B. B., van der Plas, S., Wright, K., Xu, M., Yin, H., Levell, J., and Pagliarini, R. A. (2017) Allosteric mutant IDH1 inhibitors reveal mechanisms for IDH1 mutant and isoform selectivity. Structure 25, 506–513

81. Rohle, D., Popovici-Muller, J., Palaskas, N., Turcan, S., Grommes, C., Campos, C., Tsoi, J., Clark, O., Oldrini, B., Komisopoulou, E., Kunii, K., Pedraza, A., Schalm, S., Silverman, L., Miller, A., Wang, F., Yang, H., Chen, Y., Kernytsky, A., Rosenblum, M. K., Liu, W., Biller, S. A., Su, S. M., Brennan, C. W., Chan, T. A., Graeber, T. G., Yen, K. E., and Mellinghoff, I. K. (2013) An inhibitor of mutant IDH1 delays growth and promotes differentiation of glioma cells. Science 340, 626–630

82. Chambers, J. M., Miller, W., Quichocho, G., Upadhye, V., Matteo, D. A., Bobkov, A. A., Sohl, C. D., and Schiffer, J. M. (2020) Water networks and correlated motions in mutant isocitrate dehydrogenase 1 (IDH1) are critical for allosteric inhibitor binding and activity. Biochemistry 59, 479–490

83. Albe, K. R., Butler, M. H., and Wright, B. E. (1990) Cellular concentrations of enzymes and their substrates. J Theor Biol 143, 163–195

84. Bevensee, M. O., and Boron, W. F. (2008) Effects of acute hypoxia on intracellular-pH regulation in astrocytes cultured from rat hippocampus. Brain Res 1193, 143–152

85. Bright, C. M., and Ellis, D. (1994) Hypoxia-induced intracellular acidification in isolated sheep heart Purkinje fibres and the effects of temperature. J Mol Cell Cardiol 26, 463–469

86. Bright, C. M., and Ellis, D. (1992) Intracellular pH changes induced by hypoxia and anoxia in isolated sheep heart Purkinje fibres. Exp Physiol 77, 165–175

87. Mescam, M., Vinnakota, K. C., and Beard, D. A. (2011) Identification of the catalytic mechanism and estimation of kinetic parameters for fumarase. J Biol Chem 286, 21100–21109

88. Alberty, R. A., Massey, V., Frieden, C., and Fuhlbrigge, A. R. (1954) Studies of the enzyme fumarase. III.1 The dependence of the kinetic constants at 25° upon the concentration and pH of phosphate buffers. J Am Chem Soc 76, 2485–2493

89. Persat, A., Chambers, R. D., and Santiago, J. G. (2009) Basic principles of electrolyte chemistry for microfluidic electrokinetics. Part I: Acid-base equilibria and pH buffers. Lab Chip 9, 2437–2453

90. Colman, R. F. (1975) Mechanisms for the oxidative decarboxylation of isocitrate: implications for control. Adv Enzyme Regul 13, 413–433

91. Hurley, J. H., Dean, A. M., Koshland, D. E., Jr., and Stroud, R. M. (1991) Catalytic mechanism of NADP(+)-dependent isocitrate dehydrogenase: implications from the structures of magnesium-isocitrate and NADP+ complexes. Biochemistry 30, 8671–8678

92. Ehrlich, R. S., and Colman, R. F. (1987) Ionization of isocitrate bound to pig heart NADP+-dependent isocitrate dehydrogenase: 13C NMR study of substrate binding. Biochemistry 26, 3461–3466

93. Ehrlich, R. S., and Colman, R. F. (1976) Influence of substrates and coenzymes on the role of manganous ion in reactions catalyzed by pig heart triphosphopyridine nucleotide-dependent isocitrate dehydrogenase. Biochemistry 15, 4034–4041

94. Colman, R. F. (1972) Role of metal ions in reactions catalyzed by pig heart triphosphopyridine nucleotide-dependent isocitrate dehydrogenase. II. Effect on catalytic properties and reactivity of amino acid residues. J Biol Chem 247, 215–223

95. Bacon, C. R., Bednar, R. A., and Colman, R. F. (1981) The reaction of 4-iodoacetamidosalicylic acid with TPN-dependent isocitrate dehydrogenase from pig heart. J Biol Chem 256, 6593–6599

96. Farrell, H. M., Jr., Deeney, J. T., Hild, E. K., and Kumosinski, T. F. (1990) Stopped flow and steady state kinetic studies of the effects of metabolites on the soluble form of NADP+:isocitrate dehydrogenase. J Biol Chem 265, 17637–17643

97. Dean, A. M., and Koshland, D. E., Jr. (1993) Kinetic mechanism of Escherichia coli isocitrate dehydrogenase. Biochemistry 32, 9302–9309

98. Fierke, C. A., Johnson, K. A., and Benkovic, S. J. (1987) Construction and evaluation of the kinetic scheme associated with dihydrofolate reductase from Escherichia coli. Biochemistry 26, 4085–4092

99. Miller, G. P., Wahnon, D. C., and Benkovic, S. J. (2001) Interloop contacts modulate ligand cycling during catalysis by Escherichia coli dihydrofolate reductase. Biochemistry 40, 867–875

100. Miller, G. P., and Benkovic, S. J. (1998) Deletion of a highly motional residue affects formation of the Michaelis complex for Escherichia coli dihydrofolate reductase. Biochemistry 37, 6327–6335

101. Miller, G. P., and Benkovic, S. J. (1998) Strength of an interloop hydrogen bond determines the kinetic pathway in catalysis by Escherichia coli dihydrofolate reductase. Biochemistry 37, 6336–6342

102. Miller, G. P., and Benkovic, S. J. (1998) Stretching exercises--flexibility in dihydrofolate reductase catalysis. Chem Biol 5, R105–113

103. Yen, K., Travins, J., Wang, F., David, M. D., Artin, E., Straley, K., Padyana, A., Gross, S., DeLaBarre, B., Tobin, E., Chen, Y., Nagaraja, R., Choe, S., Jin, L., Konteatis, Z., Cianchetta, G., Saunders, J. O., Salituro, F. G., Quivoron, C., Opolon, P., Bawa, O., Saada, V., Paci, A., Broutin, S., Bernard, O. A., de Botton, S., Marteyn, B. S., Pilichowska, M., Xu, Y., Fang, C., Jiang, F., Wei, W., Jin, S., Silverman, L., Liu, W., Yang, H., Dang, L., Dorsch, M., Penard-Lacronique, V., Biller, S. A., and Su, S. M. (2017) AG-221, a first-in-class therapy targeting acute myeloid leukemia harboring oncogenic IDH2 mutations. Cancer Discov 7, 478–493

104. Intlekofer, A. M., Shih, A. H., Wang, B., Nazir, A., Rustenburg, A. S., Albanese, S. K., Patel, M., Famulare, C., Correa, F. M., Takemoto, N., Durani, V., Liu, H., Taylor, J., Farnoud, N., Papaemmanuil, E., Cross, J. R., Tallman, M. S., Arcila, M. E., Roshal, M., Petsko, G. A., Wu, B., Choe, S., Konteatis, Z. D., Biller, S. A., Chodera, J. D., Thompson, C. B., Levine, R. L., and Stein, E. M. (2018) Acquired resistance to IDH inhibition through trans or cis dimer-interface mutations. Nature 559, 125–129

105. Sahai, A., Laughrey, E., and Tannen, R. L. (1990) Relationship between intracellular pH and ammonia metabolism in LLC-PK1 cells. Am J Physiol 258, F103–108

106. Qi, F., Pradhan, R. K., Dash, R. K., and Beard, D. A. (2011) Detailed kinetics and regulation of mammalian 2-oxoglutarate dehydrogenase. BMC Biochem 12, 53

107. Badur, M. G., Muthusamy, T., Parker, S. J., Ma, S., McBrayer, S. K., Cordes, T., Magana, J. H., Guan, K. L., and Metallo, C. M. (2018) Oncogenic R132 IDH1 mutations limit NADPH for de novo lipogenesis through (D)2-hydroxyglutarate production in fibrosarcoma cells. Cell Rep 25, 1018-1026.e1014

108. Marino, M. L., Fais, S., Djavaheri-Mergny, M., Villa, A., Meschini, S., Lozupone, F., Venturi, G., Della Mina, P., Pattingre, S., Rivoltini, L., Codogno, P., and De Milito, A. (2010) Proton pump inhibition induces autophagy as a survival mechanism following oxidative stress in human melanoma cells. Cell Death Dis 1, e87

109. Ananieva, E. A., and Wilkinson, A. C. (2018) Branched-chain amino acid metabolism in cancer. Curr Opin Clin Nutr Metab Care 21, 64–70

110. Avellaneda Matteo, D., Grunseth, A. J., Gonzalez, E. R., Anselmo, S. L., Kennedy, M. A., Moman, P., Scott, D. A., Hoang, A., and Sohl, C. D. (2017) Molecular mechanisms of isocitrate dehydrogenase 1 (IDH1) mutations identified in tumors: The role of size and hydrophobicity at residue 132 on catalytic efficiency. J Biol Chem 292, 7971–7983

111. Orwig, S. D., and Lieberman, R. L. (2011) Biophysical characterization of the olfactomedin domain of myocilin, an extracellular matrix protein implicated in inherited forms of glaucoma. PLoS One 6, e16347

112. Emsley, P., and Cowtan, K. (2004) Coot: model-building tools for molecular graphics. Acta Crystallogr Sect D-Biol Crystallogr 60, 2126–2132

113. Adams, P. D., Afonine, P. V., Bunkoczi, G., Chen, V. B., Davis, I. W., Echols, N., Headd, J. J., Hung, L. W., Kapral, G. J., Grosse-Kunstleve, R. W., McCoy, A. J., Moriarty, N. W., Oeffner, R., Read, R. J., Richardson, D. C., Richardson, J. S., Terwilliger, T. C., and Zwart, P. H. (2010) PHENIX: a comprehensive Python-based system for macromolecular structure solution. Acta Crystallogr Sect D-Biol Crystallogr 66, 213–221

114. Ratnikov, B., Aza-Blanc, P., Ronai, Z. A., Smith, J. W., Osterman, A. L., and Scott, D. A. (2015) Glutamate and asparagine cataplerosis underlie glutamine addiction in melanoma. Oncotarget 6, 7379–7389

115. Merk, A., Bartesaghi, A., Banerjee, S., Falconieri, V., Rao, P., Davis, M. I., Pragani, R., Boxer, M. B., Earl, L. A., Milne, J. L., and Subramaniam, S. (2016) Breaking Cryo-EM resolution barriers to facilitate drug discovery. Cell 165, 1698–1707

116. Avellaneda Matteo, D., Wells, G. A., Luna, L. A., Grunseth, A. J., Zagnitko, O., Scott, D. A., Hoang, A., Luthra, A., Swairjo, M. A., Schiffer, J. M., and Sohl, C. D. (2018) Inhibitor potency varies widely among tumor-relevant human isocitrate dehydrogenase 1 mutants. Biochem J 475, 3221–3238

